# The molecular basis of sodium-dependent fluoride export by the eukaryotic fluoride channel FEX

**DOI:** 10.1101/2025.06.25.661560

**Authors:** Chia-Yu Kang, Minjun An, Sahar Heidari, Hedieh Torabifard, Melanie D. Ohi, Randy B. Stockbridge

## Abstract

Much of life on Earth, including plants, fungi, and bacteria, evolved to resist toxic environmental fluoride. In eukaryotes, the major resistance mechanism is fluoride export by FEX proteins. Using electrophysiology and transport assays, we establish that FEX from plants and yeasts are fluoride channels whose activity depends on reversible sodium ion binding. A cryo-EM structure of FEX from *Candida albicans*, together with mutagenesis studies, reveals a fluoride permeation route through a single phenylalanine-lined pore. Molecular dynamics simulations demonstrate that a cation binding motif adjacent to the pore provides a stable sodium binding site that is accessible from the external aqueous solution. We propose that sodium gating resolves a major conundrum of channel-based fluoride efflux by preventing fluoride permeation under conditions of membrane depolarization. Comparison to bacterial fluoride channels (Flucs) provides a unique glimpse of the evolution of structural and mechanistic complexity in a membrane protein family with inverted repeat architecture.

## Introduction

Fluoride (F^-^) is present at biologically relevant concentrations throughout the natural environment, including soil, water, and certain minerals(*1*). Human use of fluoride and fluorine-containing molecules has further compounded fluoride levels in some niches(*2*). Intracellular fluoride accumulation inhibits multiple enzymes that are essential in multiple biological pathways, including glycolysis, ATP synthesis, phosphorylation, nucleotide synthesis, and other processes catalyzed by metalloenzymes (*3–5*). As a result, most microorganisms, fungi, and plants possess mechanisms to avoid fluoride toxicity(*6, 7*). Understanding these fluoride resistance mechanisms is of interest to multiple fields, including biotechnology and synthetic biology, bioremediation, agriculture, and antimicrobial development(*2, 8, 9*).

The major mechanism that plants and microorganisms use to protect against cytoplasmic fluoride accumulation is membrane export, and at least three types of membrane protein, including both antiporters and ion channels, have evolved to export this toxic halide(*10*). Bacteria possess either F^-^/H^+^ antiporters from the CLC family of anion channels and transporters known as CLC^F^(*11*), or architecturally distinct fluoride channels known as Flucs(*12*). In eukaryotes, including plants, fungi, and even some aquatic filter feeding animals, fluoride export is managed by proteins known as FEX(*7, 13, 14*). FEX possess the same fold as the Flucs, but are more architecturally complex. Whereas Flucs assemble as rare dual topology antiparallel homodimers, with one subunit inserted into the membrane facing out, and the other facing in(*15*), the FEX proteins have an inverted repeat architecture, in which their two homologous domains are synthesized as a single polypeptide with a transmembrane linker that enforces antiparallel assembly(*16*). The antiparallel architecture of the Flucs represents an evolutionary antecedent of the FEX(*17*). Although inverted repeat architecture is extremely common among membrane transport proteins(*18*), Fluc/FEX is the only family of known function in which both the antiparallel homodimer and fused inverted repeat versions are extant (*19*). This provides a unique opportunity to understand how evolutionary forces shape membrane proteins after domain duplications and fusions, which occurred in the evolutionary history of a large proportion of membrane transporters(*18*).

Functional, structural, and computational studies have yielded a detailed understanding of the architecture and fluoride transport mechanism of the Flucs. As a consequence of their antiparallel homodimeric architecture, the Flucs possess perfect two-fold symmetry(*20–22*) and a pair of antiparallel, but otherwise identical pores(*22*). Both pores simultaneously conduct fluoride, relying on the cell’s membrane potential to expel these anions (*23*). Fluoride ions enter through Fluc’s large, electropositive aqueous vestibules, and are then translocated into the dehydrated part of the permeation pathway(*24, 25*). Here, the fluorides are coordinated by a track of polar residues, as well as the electropositive ring edges of a pair of phenylalanines from a motif known as the phenylalanine box(*22, 26*). The fluorides transit rapidly through this pore, with a rate of ∼10^6^ ions per second(*12*). Anion selectivity occurs at two points along the pathway: at a conserved serine that translocates the fluoride ions from the vestibule to the constricted part of the pore(*24, 27*), and, at the opposite end of the pathway, by a gate comprised of a threonine, glutamate, and tyrosine contributed by each of three different helices(*24, 25*). Both the vestibule serine and the gating triad residues are highly conserved in prokaryotes and eukaryotes. The Flucs also possess a sodium ion buried at the dimer interface. In structures, this sodium is bound by just four ligands, all of which are main chain carbonyls from the breaks in symmetry-related transmembrane helices(*22*). This sodium is not exchangeable on biological timescales in wildtype (WT) Fluc (*22*), but has been shown to contribute to channel assembly(*28*) and is required for activity (*29*). The idiosyncratic architecture of the Flucs raises several questions about their structural relatives, the FEX. Is the symmetry and two-pore architecture of the Flucs preserved in the FEX? Are the channel mechanism and major chemical determinants of fluoride coordination retained? Does sodium or another cation play a role in the transport mechanism?

Prior functional studies, primarily with FEX homologues from *Saccharomyces cerevisiae* and *Arabidopsis thaliana*, have helped narrow these knowledge gaps, but open questions remain. Whole-cell patch recording in yeast supported functional assignment of the FEX as ion channels (*30, 31*), but did not definitively rule out other mechanisms, for example electrogenic proton-coupled transport, which has also been proposed(*32*). Sequence analysis and mutational studies suggest that the two domains of FEX are functionally asymmetric, with one functional and one vestigial pore (*13, 16, 22*). However, even for the functional pore, some residues that are critical for Fluc activity cause only mild defects when mutated in FEX. For example, a key serine responsible for ion translocation from the vestibule to the pore in Fluc does not appear to be essential for FEX-mediated fluoride resistance (*13, 24, 27*). The fluoride ion’s route through FEX thus remains uncertain. Finally, whether sodium plays a role in fluoride transport by FEX has not yet been explored.

Here we use planar lipid bilayer electrophysiology, liposome transport assays, yeast resistance assays, single-particle cryo-electron microscopy (cryo-EM), and computational simulations to establish the molecular basis of fluoride transport by FEX from plants and yeast. We find that these single-barreled electrodiffusive channels require sodium for activity, which binds reversibly to the WT channels through the external vestibule. A structure of FEX from the pathogenic yeast, *Candida albicans*, together with site directed mutagenesis, reveals a single narrow pore, lined by numerous phenylalanine residues, that bypasses the prominent vestibules characteristic of the Flucs. Together, these results provide molecular insight into fluoride transport and resistance in eukaryotes, and reveal an unprecedented glimpse into the evolutionary trajectory of a membrane protein from its most primitive dual topology form to complex inverted repeat architecture.

## Results

### Functional characterization of FEX homologs

We first surveyed genomes across the eukaryotic tree of life for FEX homologues, and found that most major eukaryotic clades include organisms that possess FEX-encoding genes (**Figure 1A**). We screened multiple phylogenetically diverse FEX homologs for expression and biochemical behavior, achieving acceptable purifications of FEX proteins from plants *A. thaliana* (FEX-AT) and tea plant *Camellia sinensis* (FEX-CS), a globally important crop plant for which fluoride resistance is highly relevant(*33*), and from yeasts *Candida albicans* (FEX-CA) and *S. cerevisiae* (FEX-SC) (**Supplementary Figure 1**). All of these FEX homologues catalyzed rapid fluoride efflux from proteoliposomes (**Figure 1B; Supplementary Figure 2A**), consistent with previous results(*14*). Similar to the prokaryotic Fluc channels, the rate of fluoride transport by the plant FEX exceeds the response time of the electrode, equivalent to a unitary turnover of >10^4^ ions/s. In contrast, the yeast homologues exhibited slower transport kinetics, with rates of 2910 ± 110 s^-1^ for *S. cerevisiae* FEX and 1050 ± 90 s^-1^ for *C. albicans* FEX (**Figure 1B**).

**Figure 1.**
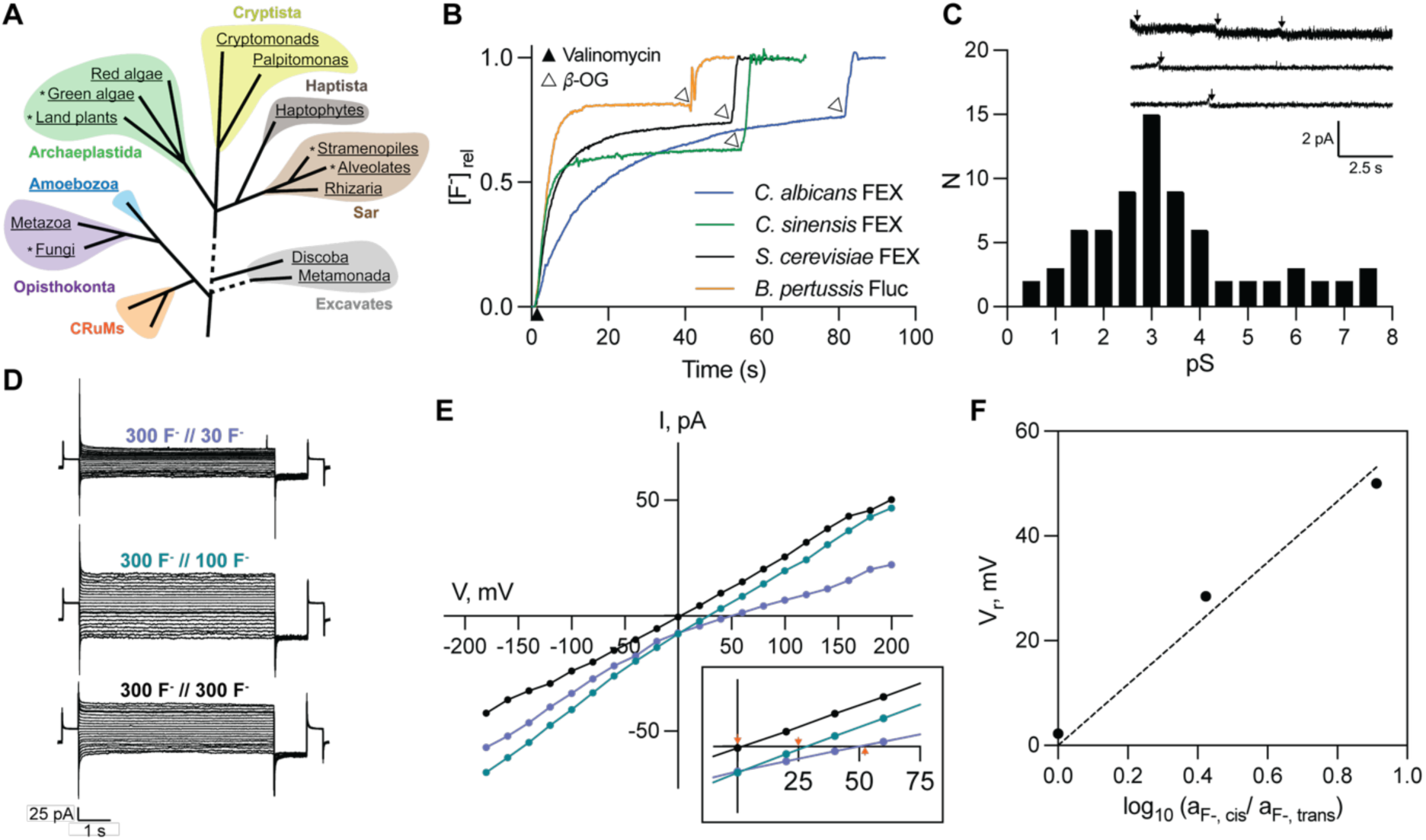
Eukaryotic FEX are fluoride channels with variable rates. (A) Eukaryotic tree of life adapted from (*80*) and annotated according to FEX distribution. Clades with species that possess FEX homologues are underlined, and clades in which FEX homologues are widespread among most species are denoted with an asterisk. FEX distribution was analyzed using UniProtKB(*81*). (B) Fluoride efflux from proteoliposomes measured with a fluoride-selective electrode. Efflux was initiated by the addition of valinomycin (closed triangle), and N-octyl-*β*-D-glucoside was added to solubilize liposomes at the end of the timecourse (open triangle). Traces are normalized against the total encapsulated F^-^. Traces are representative of 3-6 measurements. (C) Histogram of single channel insertions for FEX-CS. Representative recordings of FEX-CS as an inset, recorded at −200 mV under F^-^ gradient conditions (300 mM NaF // 30 mM NaF). (D) Representative raw current responses to families of 10-s command voltage pulses (−200 to +200 mV in 20 mV increments) from a holding potential of V=0, followed by a 500-ms tail pulse to V=-200 mV. (E) Macroscopic current-voltage (I-V) curves for three F^-^-gradient conditions at the end of the voltage pulses shown in panel D. The inset shows the I-V curve near the reversal potential, with the expected Nernstian reversal potentials for each ionic gradient indicated by the orange arrow. (F) Reversal potential (V_r_) as a function of F^-^ activity (a_F_) from three independent bilayers. Points represent the mean and SEM (error bars are smaller than the diameter of the point). The dashed line represents the expected relationship between V_r_ and the activity gradient for a selective, electrodiffusive fluoride ion channel. F^-^ activity is calculated from tabulated mean ionic activity coefficients of NaF solutions.

The rapid fluoride transport by FEX-CS enabled planar bilayer electrophysiology with the purified proteins. For liposomes reconstituted at a low protein:lipid ratio, such that each liposome is expected to bear 1-2 FEX proteins, we observe fusion to the bilayer in extremely small, but discrete steps (**Figure 1C**). Like the Flucs, the single channel currents do not exhibit sub-states or closures, and are fully open at potentials from −200 to +200 mV. A histogram of single insertion events from different bilayers yields a mean channel conductance of 3 pS (**Figure 1C**). After several dozen channel fusions, we measured the current-voltage (I-V) relationship for FEX-CS under three different fluoride gradients (**Figure 1D, E**). The measured reversal potentials were Nernstian with respect to the fluoride activity gradient (**Figure 1F**). This experiment provides unequivocal evidence of an electrodiffusive ion transport mechanism. FEX-AT behaved similarly to FEX-CS, with a 2.5 pS mean conductance and reversal potentials consistent with channel activity under fluoride gradient conditions (**Supplementary Figure 2B-D**). These experiments confirm that eukaryotic FEX acts via a channel mechanism, with substantial differences in the ion transport rates among different organisms.

### Cryo-EM structure of FEX from Candida albicans

We next sought to determine a structure of a FEX homologue using cryo-EM. At just ∼40 kDa, and without any extramembrane domains, FEX channels are a challenging target for cryo-EM analysis. To increase the particle size and aid particle alignment, we appended the 17-residue membrane-proximal external region (MPER) epitope as a continuous extension of the N-terminal transmembrane helix of FEX-CA. This epitope, derived from the envelope glycoprotein gp41 in human immunodeficiency virus (HIV), has been shown to provide a rigid, membrane-adjacent docking site for several well-characterized monoclonal antigen-binding (Fab) fragments(*34–37*) that can serve as cryo-EM fiducials for membrane proteins during single particle cryo-EM analysis(*38, 39*). In addition, the first 75 residues of FEX-CA, which are not conserved and are predicted to be disordered, were truncated to decrease structural heterogeneity. This engineered FEX-CA, with the MPER epitope, is biochemically stable and maintains comparable activity to the full-length FEX-CA (**Figure 2A and Supplementary Figure 1A, D**). The Fab 10E8v4 forms a stable complex with MPER-tagged FEX that is monodisperse by size-exclusion chromatography (SEC) (**Figure 2B**) with homogenous particles (**Supplementary Figure 3A**). Binding of this Fab has no effect on the fluoride efflux activity of FEX-CA (**Figure 2A**).

**Figure 2.**
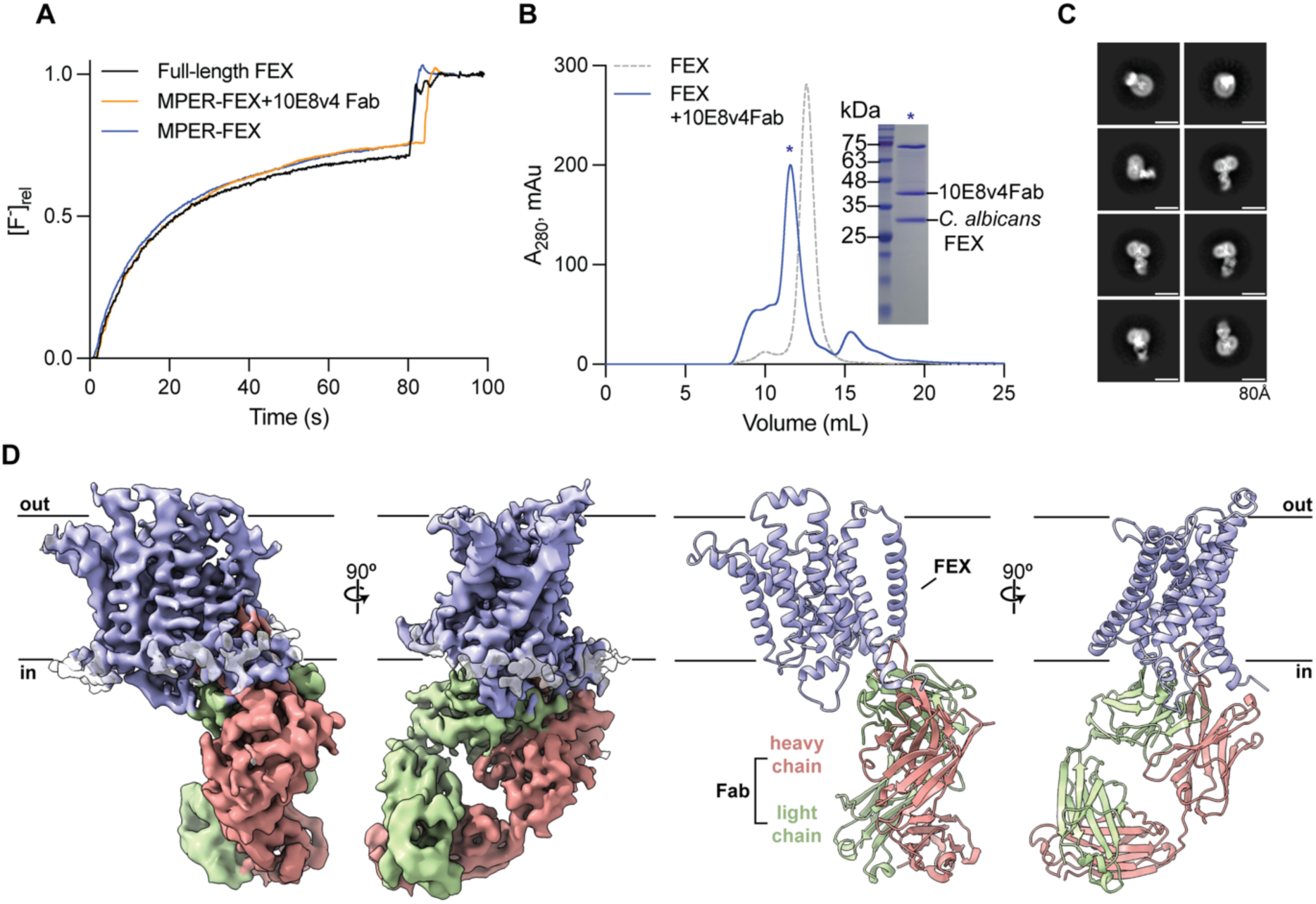
Epitope tag, antibody complexation and structure determination for *C. albicans* FEX. (A) Fluoride efflux from proteoliposomes reconstituted with the indicated constructs. Traces are normalized against total encapsulated F^-^. Traces are representative of 3-6 measurements. (B) Size exclusion chromatograms of MPER-tagged FEX-CA alone (gray dashed line) or with 10E8v4 Fab at a 1:1 molar ratio (blue trace). Inset: Coomassie-stained SDS-PAGE of the peak fraction for the FEX-CA/10E8v4 trace (marked by the asterisk in the chromatogram). (C) Representative 2D averages of 10E8v4 Fab-bound FEX-CA in DDM micelles, with an 80 Å scale bar indicated. (D) Left, cryo-EM map of FEX-CA/10E8v4 Fab complex. Right, model of FEX-CA/10E8v4 Fab complex. For both panels, FEX is shown in purple, the 10E8v4 heavy chain in salmon pink, the 10E8v4 light chain in green. The lines represent the approximate boundaries of the plasma membrane with the cytoplasmic side (in) and extracellular side (out) labeled.

The Fab-bound MPER-FEX-CA, solubilized in DDM micelles, was used for single particle cryo-EM analysis. Selected 2D averages of Fab-bound proteins in different orientations exhibit transmembrane domain features (**Figure 2C**). After multiple rounds of ab-initio 3D reconstruction, 3D classification, heterogenous refinement, and non-uniform refinement, we obtained a 4.0 Å density map of MPER-FEX-CA bound to the 10E8v4 Fab, with local resolution up to 3.5Å in the protein core (**Figure 2D; Supplementary Figure 3; Supplementary Figure 4**; (**Supplementary Table 1**). In this map, the densities for the constant and variable domains of the 10E8v4 Fab are well resolved, and in FEX-CA, all nine transmembrane (TM) α-helices and their connecting loops have clearly resolved sidechains sufficient for model building and residue assignment (**Supplementary Figure 5**). Docking the known 10E8v4 structure (PDB 5IQ9)(*37*) into the map shows that the Fab fragment’s interactions with FEX-CA are almost entirely restricted to the MPER epitope, avoiding perturbations of mechanistically important parts of the protein, a common problem for fiducials selected from randomized libraries(*40*). We built an atomic model for residues 1-381 of the engineered FEX-CA construct, corresponding to 76-381 of WT FEX-CA (**Supplementary Table 1**). Only nine residues from the unstructured C-terminus were missing from the density map and could not be modelled. This structure shows that FEX-CA has homologous N-terminal (TM 1-4) and C-terminal domains (TM 6-9), which comprise the protein core. The domains are connected by a transmembrane linker helix (TM 5) that maintains the antiparallel domain architecture (**Figure 3A**). As in the Flucs, the third helix in each domain (TM3 and TM8) is broken by a stretch of approximately three non-helical residues. These breaks in the α-helices cross over each other at approximately the midpoint of the membrane. Compared to our experimental structure, the model predicted by AlphaFold 3(*41*) has a root mean squared deviation (RMSD) of 2.5 Å for 237 C_⍺_ atoms within the transmembrane helices. While the protein core (N- and C-terminal domains) of the experimental and Alphafold models are similar (RMSD of 1.6 Å for 213 C_⍺_ atoms in the transmembrane helices), significant deviations occur in the vicinity of TM5, the linker helix (**Supplementary Figure 6)**.

**Figure 3.**
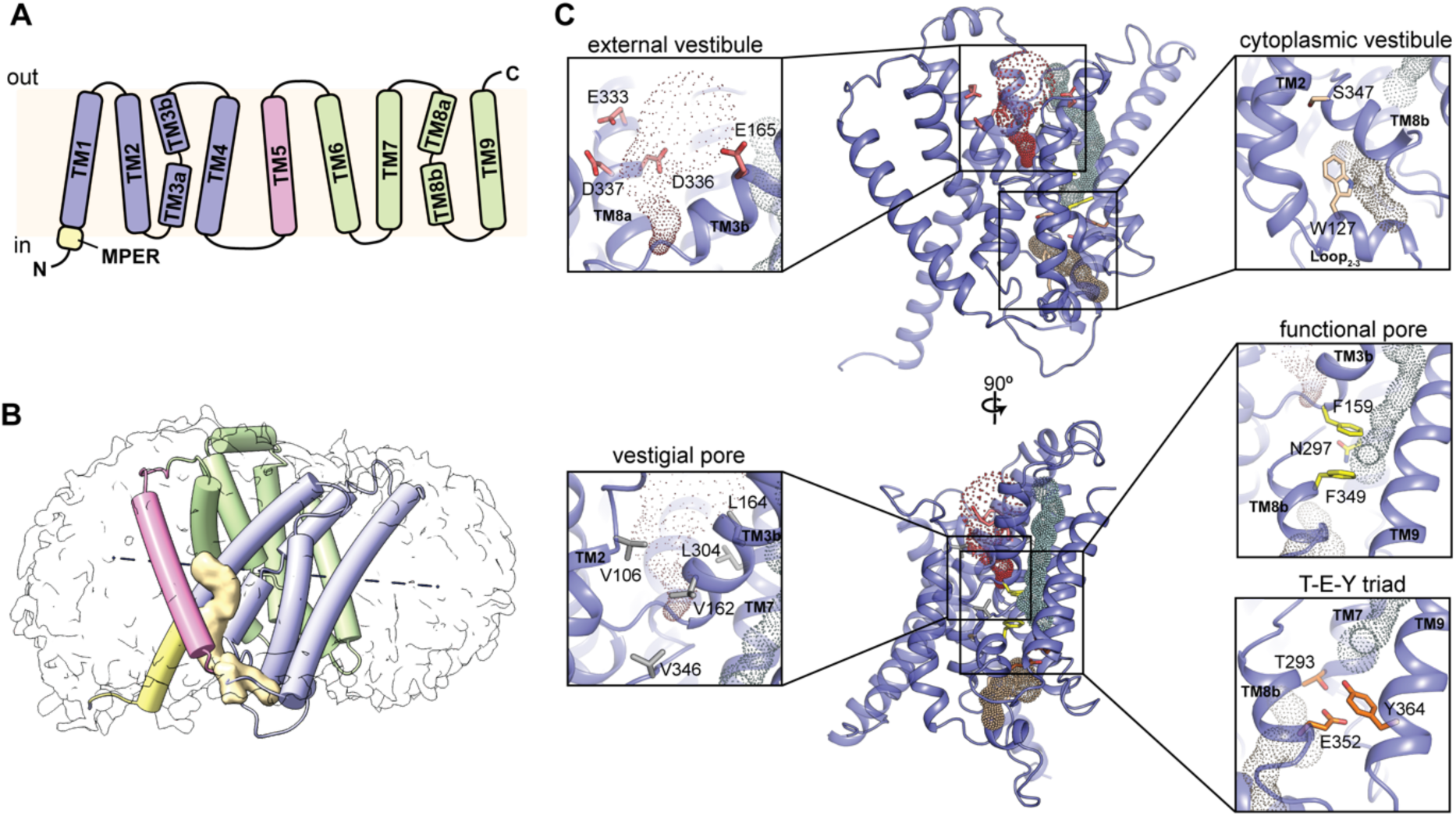
Domain duplication and architectural asymmetry in FEX. (A) Transmembrane topology of MPER-tagged FEX-CA. (B) Helical model of FEX-CA with coloring as in panel A: pseudosymmetric domains colored in green and purple, TM5 linker helix in pink, and MPER tag in bright yellow. The micelle density is shown in gray. The internal pseudosymmetry axis is shown as a dashed line, and the map for a detergent-like non-protein density that segregates the linker from the core is shown in light yellow. (C) Model of FEX-CA with aqueous cavities determined using CAVER(*82*) shown as dots and conserved residues indicated and shown as sticks. Insets show zoomed in views of key structural features.

### Architectural asymmetry of FEX

FEX possesses internal C2 pseudosymmetry with the N- and C-terminal domains tilted along a symmetry axis approximately 10° off the membrane normal (**Figure 3B**). This contrasts with the Flucs, which exhibit perfect 2-fold domain symmetry about an axis parallel to the plane of the membrane(*22, 40*). The N- and C-terminal domains of FEX-CA align well across TM α-helices 1-3, with an RMSD of 2.5 Å. However, rigid body rotations of fourth helix in each domain, TM4 and TM9, in relation to the rest of their domain bundle introduces a ∼20° offset in the structural alignment of the N- and C-terminal domains (**Supplementary Figure 7**). A non-protein density consistent with a lipid or a detergent molecule segregates the linker helix, TM5, from the protein core, so that the core and linker are limited to a single point of contact: one putative hydrogen bond between S235 in the linker helix and H80 in the N-terminal core domain (**Figure 3B**). These structural observations are consistent with the lack of conservation in TM5 sequences among eukaryotic FEX.

In the Fluc dimers, the crossover of the broken helices 3 and 3’ defines the channels’ most prominent feature, a pair of aqueous, electropositive vestibules. These vestibules serve as an entrypoint and electrostatic funnel for fluoride ions before they enter the constricted part of the pore (*24, 25*). The vestibule architecture is substantially different in FEX-CA. The cytoplasmic vestibule of FEX-CA is plugged by the loop between TM2 and TM3a, which forms a short helical segment and extends the bulky side chain W127 into the cleft defined by the broken helices of TM3 and TM8 (**Figure 3C**). Because of this occlusion, there is little aqueous volume in the cytoplasmic vestibule. Meanwhile, the external vestibule, while water-filled and of similar size as the vestibules in the Flucs, is strikingly electronegative. Several acidic sidechains, including E165, E333, D336, and D337, line the aqueous surface (**Figure 3C**). These electrostatics would disfavor fluoride entry into the external vestibule, which is likely to be adaptive in the presence of high environmental fluoride.

In lieu of a commodious vestibule to funnel fluoride ions into the pore, our FEX-CA model instead reveals a long, narrow aqueous crevice surrounded by TM3b, TM4, TM8b, and TM9 (**Figure 3C**). This aqueous route traverses almost the entire length of the protein, interrupted only by the conserved triad of T293, E352, and Y364. In Flucs, this triad contributes to ion selectivity(*24*); the position of these interacting residues on the cytoplasmic side of the pore in FEX is consistent with a role as the primary selectivity filter for fluoride ion.

The fluoride permeation pathway in FEX-CA partially coincides with one of the pores of the double-barreled Flucs, skirting the two conserved edges of the phenylalanine box motif (F159 and F349) and the conserved N297 near the mid-point **(Figure 3C**). Consistent with mutagenic studies(*13*), this structure shows in FEX, the second pore has degraded. The opposite two edges of the phenylalanine box motif have been replaced with a valine and an isoleucine, and the vestigial route is tightly packed with hydrophobic sidechains (**Figure 3C**).

### The fluoride permeation pathway is lined by phenylalanines and bypasses the vestibules

The aqueous crevice is lined by several polar residues expected to be involved in fluoride permeation, including residues T293, E352, and Y364 from the selectivity triad, and the polar track sidechain N297 (**Figure 4A**). These positions are highly conserved from bacteria to eukaryotes, and have been shown to be essential for fluoride resistance by *S. cerevisiae* FEX (*13, 16*). In addition to these expected polar sidechains, the FEX-CA pore is also lined by residues that have not previously been implicated in eukaryotic fluoride resistance, notably a confluence of phenylalanines surrounding the pore along its entire length (**Figure 4A**). The conserved phenylalanines from the phenylalanine box, F159 and F349, which have previously been shown to contribute to FEX function(*13*) are joined by F167, F191, F373, and F294 to line the aqueous route. Coordination by the electropositive quadrupolar edge of aromatic sidechains has emerged as a theme in biological fluoride recognition, so it seems likely that these phenylalanines also contribute to fluoride permeation(*22, 26, 42*).

**Figure 4.**
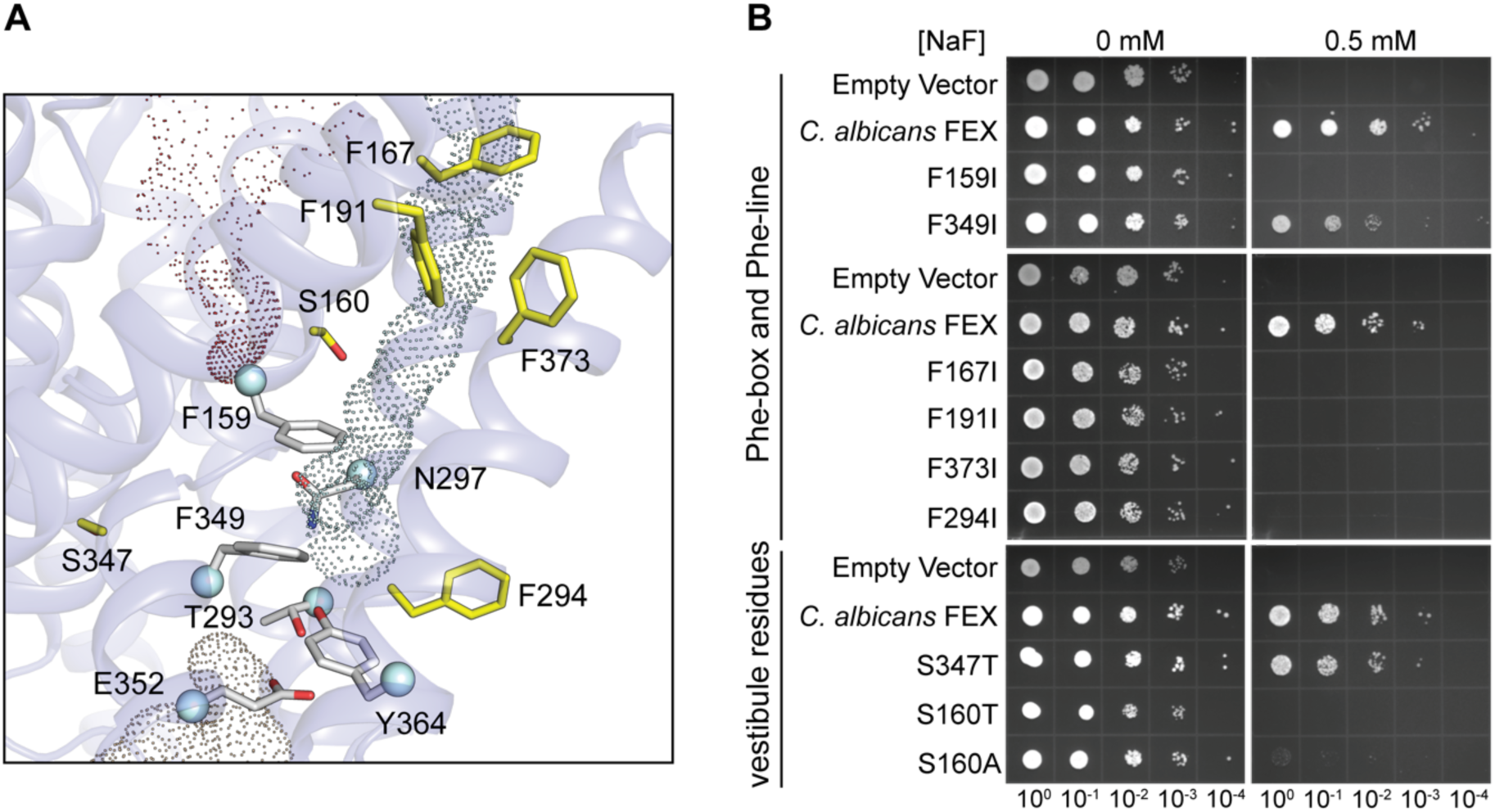
The fluoride permeation pathway of FEX-CA. (A) Zoomed in view of the predicted permeation pathway (dots; determined using CAVER(*82*)). Conserved residues previously implicated in FEX function are shown as gray sticks. Sidechain sticks for pore-lining residues newly identified in this study are shown as yellow sticks. (B) Ten-fold serial dilutions of Δ*fex1*Δ*fex2 S. cerevisiae* expressing the indicated FEX constructs on plates with fluoride as indicated. Experiments are representative of 2-3 independent biological replicates, with each replicate having 2 technical replicates.

To test this, we introduced constructs with phenylalanine-to-isoleucine point mutations into a fluoride-sensitive yeast strain (*S. cerevisiae* Δ*fex1*Δ*fex2*(*7*)) and monitored the fluoride resistance relative to cells expressing WT FEX-CA (**Figure 4B**). We selected isoleucine sidechains to maintain the hydrophobicity and size of phenylalanine, while eliminating the polarizability of the aromatic ring. Although Western blot analysis showed that the membrane expression varied as much as five-fold among these constructs (**Supplementary Figure 8B, C**), the expression level was not a decisive factor in the fluoride resistance phenotype. For example, F349I exhibited the lowest expression level among the mutants, at just 20% of WT, and yet showed the most robust fluoride resistance among the mutant constructs. F349I thus provides a benchmark for a lower limit of FEX expression that still rescues the fluoride sensitive strain.

These experiments support the hypothesis that the Phe box residues contribute to fluoride resistance: the F159I mutant did not grow at 0.5 mM NaF, whereas F349I was less impaired, only showing sensitivity at 5 mM NaF that might be explained by this construct’s lower expression level (**Figure 4B, Supplementary Figure 8A**). This is consistent with previous observations in bacteria and yeast, where mutation of the F349 equivalent is less detrimental than mutation of the F159 equivalent(*13, 22*). Mutations to the other Phe residues that line the permeation pathway, F167I, F191I, F373I, and F294I, all cause substantial fluoride sensitivity at 0.5 mM NaF (**Figure 4B**), establishing the pore-lining phenylalanines as a key component of fluoride export activity *in vivo*.

Notably, the pore crevice bypasses the cytoplasmic aqueous vestibule, which is functionally important in the Flucs. However, it is conceivable that rearrangement of the TM2-TM3 loop segment could unplug FEX-CA’s vestibule and present an alternative entry point for fluoride into the pore. We therefore sought further functional confirmation that the cytoplasmic vestibule is not part of the fluoride permeation route. To test this, we mutated S347 (**Figure 4A**), a position analogous to the vestibule serine in the Flucs that is proposed to rotamerize to deliver fluoride from the vestibule to the pore(*24, 27*). In the Flucs, mutation to the smaller residue alanine has little impact on fluoride transport, but introducing the conformationally restricted threonine at this position abolishes activity(*24*). In contrast, in FEX the analogous mutation S347T shows robust rescue of the fluoride-sensitive phenotype (**Figure 4B, Supplementary Figure 8A**). This suggests that a similar conformational change is not required of the vestibule-facing S347, and, by inference, that ion translocation from the cytoplasmic vestibule to the pore is not a feature of FEX’s transport mechanism. Other mutations introduced at this position in the *S. cerevisiae* homologue have similarly been shown to be functionally mild(*13, 16*).

Meanwhile, in the FEX-CA structure, the symmetry related serine, S160, is rotated away from the external vestibule, in an orientation similar to the alternate rotamer proposed, but not structurally verified in the Flucs(*24, 27*). The FEX-CA density map shows that the S160 sidechain interacts with the T301 sidechain from TM7, ∼5 Å away from the aqueous crevice (**Supplementary Figure 9**). In contrast to *S. cerevisiae* FEX, which is relatively insensitive to mutation at this position(*16*), in *C. albicans* FEX, S160A and S160T both greatly diminish fluoride resistance (**Figure 4B**). However, given this sidechain’s distance from both the vestibule and the aqueous crevice and its lack of functional conservation in the *S. cerevisiae* protein, we think it is unlikely that S160 is directly involved in fluoride transport by FEX.

### Sodium is essential for FEX channel activity

In the Flucs, structures show a sodium ion is buried at the dimer interface, coordinated by four carbonyl oxygen atoms from the crossover region of TM3 helical breaks (*29*). The sodium is inserted during Fluc assembly and is not exchangeable in WT Fluc(*28, 43*). It is not known whether Na^+^ or other cations play any role in FEX structure or function.

To investigate whether Na^+^ contributes to FEX activity, we examined fluoride efflux from proteoliposomes as a function of Na^+^ concentration for WT FEX-CA and FEX-CS. For both homologues, we observe a sodium dose-dependent increase in total fluoride efflux (**Figure 5A)**, reflecting a change in the fraction of liposomes that possess an active fluoride exporter. Western blot analysis of the proteoliposomes confirmed that this trend is not related to protein reconstitution efficiency (**Supplementary Figure 10**). We instead interpret this result to mean that the relative proportion of active and inactive FEX varies as a function of Na^+^ concentration. Using Poisson distribution statistics to correct for liposomes that contain multiple channels(*44*), we converted our measurement of transport-active liposomes into an estimate of the fraction of active protein at each Na^+^ concentration. This relationship is well fit by a single-site binding isotherm for both FEX-CA and FEX-CS, with dissociation constant (K_d_) values of 0.4 mM and 0.5 mM respectively (**Figure 5B**). It is notable that for both homologues, there is residual fractional activity in the absence of Na^+^, which is unexpected for an equilibrium two-state model if Na^+^ is the only cation that binds FEX and supports fluoride efflux. We therefore considered that K^+^ might substitute for Na^+^ at the high concentrations (300 mM K^+^) used for these assays. Indeed, when the background K^+^ concentration is reduced to 100 mM, the proportion of transport-inactive liposomes increases (**Supplementary Figure 11**), consistent with weak K^+^ binding supporting fluoride efflux. Li^+^, in contrast, inhibits fluoride export activity by FEX-CA and FEX-CS, which is reduced to background levels at 10 mM Li^+^(**Figure 5C**).

**Figure 5.**
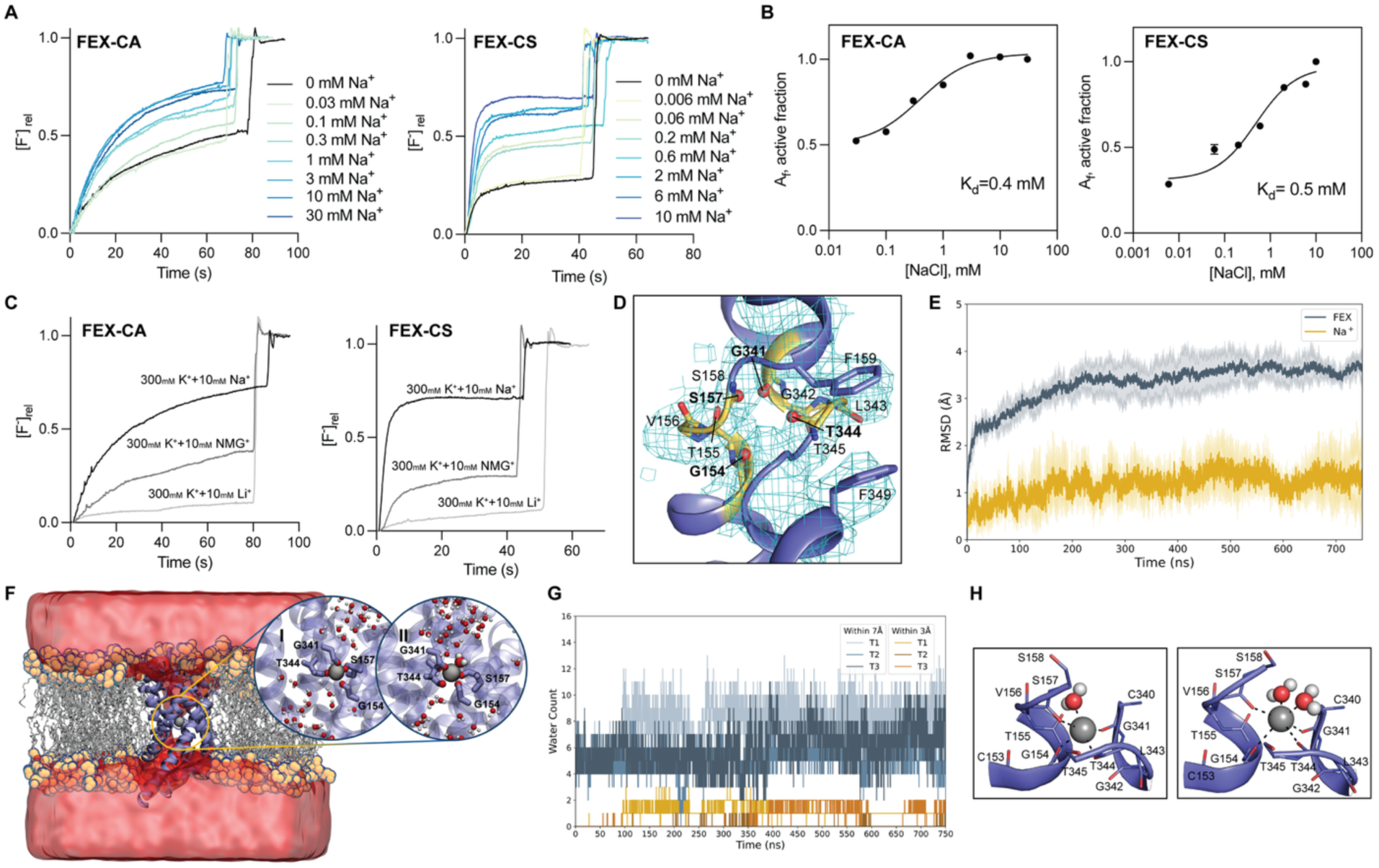
Sodium-dependent transport and binding by FEX-CA. (A) Representative traces showing fluoride efflux from proteoliposomes reconstituted with WT FEX-CA (left) or FEX-CS (right) as a function of Na^+^ concentration. Traces are normalized against the total encapsulated F^-^ determined after detergent addition. (B) The fraction of active protein (A_f_) for FEX-CA (left) and FEX-CS (right) as a function of Na^+^. The solid shows a fit to the data with a single-site binding model with K_d_ values of 0.4 mM and 0.5 mM for FEX-CA and FEX-CS, respectively. Points and errors bars represent the mean and SEM of 3 independent measurements. When not visible, error bars are smaller than the diameter of the point. (C) Fluoride efflux from FEX-CA (left) or FEX-CS (right) proteoliposomes in the presence of the indicated cations, normalized as in panel (A). Traces are representative of three independent experiments. (D) Map and model of the TM3-TM8 helical breaks. The carbonyl oxygen atoms proposed to interact with cations are shown as red spheres. The corresponding residues are shown in yellow stick representation. (E) RMSD of FEX-CA and the Na^+^ ion placed in the niche. The solid lines and shaded areas show the average and standard deviation (s.d) of three independent simulations, respectively. (F) A representation of water penetration into protein. Panel I and II show water penetration at 92 ns and 623 ns in trial 1, respectively. (G) Waters within 7 Å (blue shades) and 3 Å (yellow shades) of the Na^+^ ion as a function of simulation time. Each trial is shown in a different shade. (H) Examples of Na^+^ coordination geometry observed during simulations. The structures shown in (H) are rotated by 180° compared to panels I and II in (F).

While we do not see sodium in our map, and do not expect to at this resolution, the protein is structured to accommodate a sodium at a similar position as the Flucs. Well-defined density of the crossover region shows four carbonyl oxygens made available by the TM breaks poised near the bottom of the external vestibule, arranged in a cation-binding motif called a niche(*45*) (**Figure 5D**). These positions, G154, S157, G341, and T344, are equivalent to those that bind Na^+^ in the Flucs. The pore-lining residues F159 and F349 are immediately adjacent to the residues that comprise the niche motif, and pore-lining N297 also hydrogen bonds with the helical breaks, potentially linking cation binding and fluoride export activity.

To evaluate the stability of sodium in this putative binding site, we performed molecular dynamics simulations of FEX-CA with a Na^+^ ion initially placed in the niche. In each of 3 ∼750 ns simulations, the Na^+^ ion remained coordinated by the four carbonyl oxygens atoms at close distances for the entire simulation (**Figure 5E, Supplementary Figure 12A**). This stable coordination is supported by a strong electrostatic attraction, with a van der Waals (vdW) component consistent with a close interaction between the Na^+^ and FEX residues (**Supplementary Figure 12B**). The residues that coordinate Na^+^ ion exhibited a low root mean squared fluctuation (RMSF) relative to the rest of the protein (**Supplementary Figure 12C**).

In these simulations, water penetrates the protein, especially the external vestibule (**Figure 5F**). Approximately 5-10 waters are within 7 Å of the Na^+^ at any given time (**Figure 5G**). Water molecules from the external vestibule move in and out of the sodium ion’s coordination sphere, typically exchanging quickly with other nearby waters (**Figure 5G**). The electrostatic interaction energies indicate a favorable interaction between the Na⁺ ion and nearby water molecules. Initially, the electrostatic energy is close to zero when no water molecules are present around the sodium ion, but becomes increasingly negative as hydration occurs during the simulation (**Supplementary Figure 12D**). Most commonly, a single water coordinates Na^+^ from an axial position, yielding distorted trigonal bipyramid sodium coordination geometry, although instances of two waters binding were also observed, yielding 6-coordinate distorted octahedral geometry (**Figure 5H**). Thus, these simulations demonstrate that the cation niche provides a binding site for sodium ion that is accessible to the external vestibule.

## Discussion

A substantial proportion of membrane transport proteins possess inverted repeat architecture that evolved from the duplication and fusion of subunits that have since been lost to evolutionary history. The structural and mechanistic features of FEX that we have uncovered in these analyses can be examined through this lens (**Figure 6**). Thus, in addition to providing the first molecular blueprint for fluoride transport in eukaryotes, this study also represents a window into the evolution of macromolecular complexity and specialization in eukaryotic transport proteins with inverted repeat architecture.

**Figure 6.**
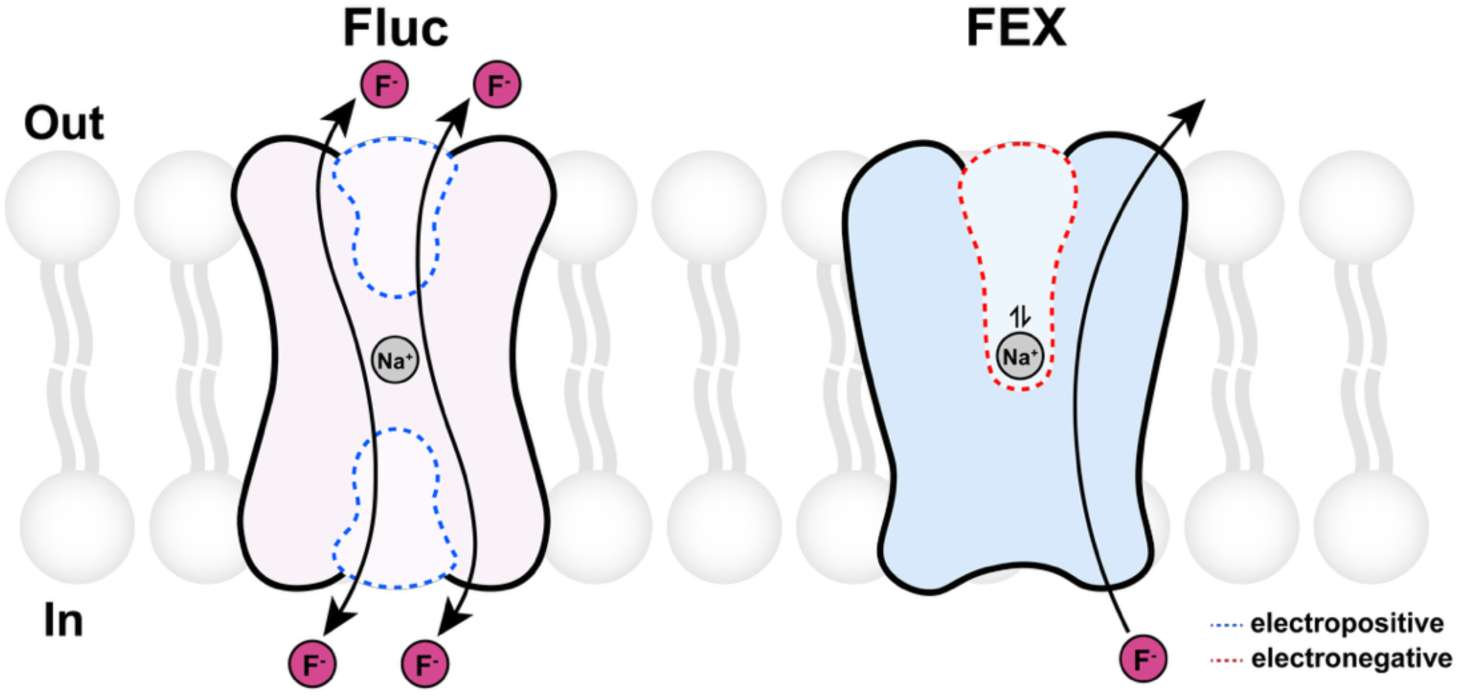
Structural and mechanistic innovations of inverted repeat FEX. Schematic of major differences between bacterial dual topology homodimer Fluc and the eukaryotic two-domain, inverted repeat FEX. Differences depicted include (1) two pore architecture in Fluc versus single pore architecture in FEX, (2) changes to vestibule architecture, electrostatics and the involvement of the vestibules in the permeation pathway, (3) a buried Na^+^ at the dimer interface in Fluc versus reversible Na^+^ binding that modulates channel function in FEX. We propose that, in FEX, the Na^+^-dependent activity guards against fluoride influx. Vestibules are indicated by dashed lines in blue (electropositive) or red (electronegative).

Our data support four main structural and mechanistic insights: First, the diffusive mechanism of fluoride transport is preserved, but different eukaryotic homologues possess rates that differ over many orders of magnitude, implying that rapid export is not physiologically necessary in all circumstances. Second, features that are functionally redundant in Fluc have degraded in FEX. In particular, the entire second pore is absent in the eukaryotic fluoride channels. The pore that is preserved in FEX maintains many of the key residues that enable fluoride transport in Flucs, and also incorporates several new phenylalanine sidechains along the length of the aqueous route. Third, relief of the symmetric constraint of the Flucs allowed specialization in FEX, such as the increased electronegativity of external vestibule, which we hypothesize prevents anion accumulation near the external face. And fourth, the sodium ion, which is purely structural in bacterial Flucs, is exchangeable in WT FEX, and we speculate that this is functionally important under physiological scenarios.

Regarding the first of these key differences, the rate of transport, it has never been obvious that fluoride resistance demands the rapid efflux (10^6^ ions per second) characteristic of the Fluc proteins, particularly because many bacteria instead rely on CLC^F^ antiporters to expel fluoride ions. CLC^F^s have subdued transport rates of 100-1000 fluoride ions/second(*11, 46*), on par with FEX-CA, the slowest FEX homologue examined in this work. In accord with its comparatively slow transport rate, FEX-CA lacks features that are associated with fast ionic flux in the Flucs and other ion channels, such as an electrostatically complementary vestibule that concentrates anions within the protein. Instead, the cytoplasmic vestibule of FEX-CA is occluded by an α-helical segment of the TM2/3 linker. The length and composition of the TM2/3 linker varies considerably among different FEX homologues (**Supplementary Figure 13**). In the plant proteins, FEX-CS and FEX-AT, which exhibit currents sufficiently large for detection by single molecule electrophysiology, this linker is shorter and lacks the tryptophan that extends most deeply into the vestibular volume. We speculate that homologue-specific properties of the TM2/3 linker contribute to the rate of fluoride export exhibited by different FEX homologues.

The second key insight from our structural and functional studies is the remodeling of the pores in FEX relative to Fluc. Given the functional redundancy of Fluc’s paired pores, evolutionary theory predicts that non-functionalization of one of the two pores is likely(*47*). This outcome has long been suspected from sequence analysis and mutagenesis(*13, 22*). Our current work shows how hydrophobic sidechains have come to pack the pore in the N-terminal domain, occluding the aqueous route. Some sidechains from this vestigial pore have nonetheless been maintained by evolution, including R89, N110, and S160(*13, 16*). For all of these, our structure reveals hydrogen bonding interactions with other protein sidechains or main chain atoms, especially from the crossover region, highlighting an ancillary role in maintaining protein structure for these conserved residues.

An unanticipated feature of the FEX-CA structure was the degree to which FEX’s remaining functional pore has been re-routed through the protein compared to Fluc. The occlusion of the vestibule that we observe in the structure, and the preservation of fluoride transport by key vestibule mutant S347T both suggest that fluoride does not enter the protein via this vestibule in FEX. Without this deep entry point, (in Fluc the vestibule accounts for nearly half the total membrane span that fluoride ions must cross), the ions instead approach the conserved phenylalanine box region via a narrow crevice along TM4 that is approximately perpendicular to the membrane. This extended constriction requires several additional residues to line its full length. Accordingly, four additional phenylalanines are positioned each with its electropositive quadrupolar edge available to coordinate anions in the pore. This presents another example of the importance of polarizable aromatics to the recognition and transport of fluoride and other halides by ion channels and transporters(*22, 42, 48*).

The final evolutionary innovation of the FEX is the repurposing of the structural sodium as a functional effector. In FEX-CA the sodium binding niche motif is solvent accessible via the electronegative external vestibule, and our functional data demonstrate that the Na^+^ is exchangeable and required for full fluoride efflux activity for WT FEX-CS and FEX-CA. The apparent K_d_ for Na^+^ is in the low millimolar range, falling within a typical biologically responsive range for microorganisms – similar, for example, to several microbial Na^+^-coupled transporters(*49–51*). Moreover, we find that the connection between cation binding and channel activity is highly specific, since Li^+^ competes with Na^+^ for binding but does not support fluoride efflux, despite differing in ionic radius by just 0.26 Å.

These observations provoke the question of whether sodium binding is relevant to FEX function under physiological circumstances. We propose that the central Na^+^ solves a key conundrum of channel-based fluoride efflux, which is that fluoride expulsion requires the positive-outside potential of a polarized membrane. Membrane depolarization, which occurs under conditions of low energy balance or stress, would leave cells vulnerable to undesirable influx of environmental fluoride. We suggest that sodium binding could mechanistically link membrane depolarization and channel activity to restrict fluoride backflow during periods of low cellular energy. For a metabolizing cell, the membrane potential would drive the sodium cation into its binding site at the bottom of the external vestibule, supporting fluoride flow when the electrical gradient favors efflux. Membrane depolarization, on the other hand, would tilt the equilibrium toward cation dissociation and thus channel closure, preventing fluoride influx under these conditions. This does not have to be a fast gating mechanism, since membrane depolarizations in plants or microbes are typically long.

In summary, our findings establish a foundation for understanding fluoride physiology and transport in eukaryotes, and provide a unique perspective on the evolution of structural and mechanistic complexity in membrane transport proteins. The evolutionary arc of the Fluc/FEX family reprises various evolutionary themes, including the degradation of redundancy in the pores, the specialization of the independently evolving vestibules after duplication, and the evolutionary co-option of the structural sodium into a functional role.

## Materials and Methods

### Expression constructs and mutagenesis

Protein expression constructs (**Supplementary Table 2**) were composed of codon-optimized synthetic genes encoding FEX homologues (Integrated DNA Technologies, Coralville, IA) with the MPER tag (*38*), an N-terminal pre-signaling peptide of ⍺-mating factor to improve mature protein secretion and insertion into the *S. cerevisiae* plasma membrane(*52*), and a C-terminal streptavidin tag for protein purification. These constructs were cloned into the pDD_Leu2D vector(*53*) and transformed using a standard lithium acetate method into *S. cerevisiae* strain BJ2168 for protein expression, or Δ*fex1*Δ*fex2* SSY3(*7*) for yeast resistance assays. All constructs were verified by sequencing. Mutagenesis was performed using overlap extension PCR with primers synthesized by Integrated DNA Technologies (**Supplementary Table 3**).

### FEX expression, purification, and reconstitution

Starter cultures were prepared from yeast transformants in leucine dropout media (SC-Leu, Sunrise Science Products, Knoxville, TN) with 2% glucose, and incubated overnight at 30 °C, 240 rpm prior to inoculation of protein expression cultures in SC-Leu with 2% lactate and 0.2% glucose. These were grown (30 °C, 240 rpm) until an OD_600_ of 0.8, then induced with galactose (final concentration 2% v/v). After 24 hours, cell pellets were harvested, washed with Tris-EDTA (TE) buffer, snap-frozen in liquid nitrogen, and stored at −80°C until use.

Protoplasts were prepared from yeast pellets using Zymolase 20T digestion of the cell wall (20 U/g cell pellet) according to the manufacturer’s protocol (Amsbio, Cambridge, MA). Protoplasts were harvested by centrifugation, resuspended in breaking buffer (20 mM Tris-HCl, pH 8.0, 100 mM NaCl, 5 mM NaF), and lysed using ∼25 strokes in a pre-chilled dounce homogenizer. Lysate was extracted with 2% n-Dodecyl-β-D-Maltoside (DDM) (Anatrace, Maumee, OH) for 2 hr at room temperature. After pelleting cell debris, supernatant was applied to a gravity column with Strep-Tactin XT 4Flow resin (IBA Lifescience, Pittsburgh, PA) equilibrated with wash buffer (breaking buffer with 1 mM DDM). Protein-loaded resin was washed with 20 column volumes (CV) of wash buffer and eluted with 6 CV of wash buffer with 50 mM biotin. Eluate was further purified by size-exclusion chromatography (Superdex 200, Cytiva, Marlborough, MA) in SEC buffer (20 mM 4-(2-Hydroxyethyl)piperazine-1-ethanesulfonic acid (HEPES) pH 7.5, 150 mM NaCl, 5 mM NaF, and 0.5 mM DDM for cryo-EM or 1 mM DDM for functional studies). Fluc-Bpe was expressed and purified exactly as previously described(*22*).

Freshly purified protein was added to *E. coli* polar lipids (EPL; Avanti Polar Lipids, Alabaster, AL) solubilized with 35 mM 3-[(3-Cholamidopropyl)-Dimethylammonio]-1-Propane Sulfonate (CHAPS; Anatrace, Maumee, OH) followed by dialysis against 4-6 L of reconstitution buffer (typically 15 mM HEPES pH 7.5, 300 mM KF and 10 mM NaF) over 1.5 days. For cation-dependent experiments, NaF was substituted with NaCl, N-methyl-glucamine chloride, or LiCl as indicated. After dialysis, protein/lipid mixtures were subjected to three freeze/thaw cycles and extruded through a 400 nm filter (Cytiva, Marlborough, MA) to generate unilamellar proteoliposomes. Proteins were reconstituted at a protein:lipid ratio of 1 µg protein/mg EPL (protein:lipid molar ratio of ∼1:50000) for the FEX homologues or 0.2 µg protein/mg EPL (molar ratio of 1:200000) for Fluc-Bpe.

### Fab preparation

10E8v4 and VRC42 antibody expression vectors were obtained from the AIDS Reagent Program, Division of AIDS, NIAID, NIH from Dr. Peter Kwong (*37, 38, 54*). Heavy and light chain DNA was transfected into ExpiCHO-S and expressed using ExpiFectamine CHO reagents (Thermo Fisher Scientific) according to the manufacturer’s protocol. Six days post-transfection, supernatant was collected and the IgG-10E8v4 purified by gravity-flow affinity chromatography using protein A Sepharose 4B resin (Invitrogen) and eluted with Pierce IgG Elution buffer (Thermo Fisher Scientific). After pH neutralization, the 10E8v4 Fab fragment was generated by papain cleavage (Sigma-Aldrich 1:200 v/v) in phosphate buffered saline with 1 mM EDTA and 6.5 mM cysteine, followed by protein A affinity column purification to separate the Fc domain and Fab fragment, and SEC (Superdex 200, 20 mM HEPES pH 7.5, 150 mM NaCl, and 5 mM NaF).

### Cryo-EM grid preparation and data collection

MPER-tagged FEX-CA was incubated with the 10E8v4 Fab at a molar ratio of 1:1 for 1-2 hr at room temperature. The complex was isolated using a Superdex 200 column equilibrated in SEC buffer. This sample was concentrated to 3-4 mg/mL and 3.5 µl was applied to glow-discharged copper Quantifoil R1.2/1.3 300 mesh grids (Electron Microscopy Sciences, Hatfield, PA, USA). Grids were blotted for 2.5 s at 4 °C, 100% humidity and plunge-frozen in liquid nitrogen cooled liquid ethane using a Vitrobot Mark IV (Thermo Fisher Scientific). Grids were imaged on a Titan Krios G3 electron microscope (Thermo Fisher Scientific) operated at 300 kV and equipped with Gatan K3 direct electron detector and a Bioquantum Imaging Filter (slit width 20 eV). Micrographs were collected at magnification of ×81,000 (1.1 Å per pixel) or ×105,000 (0.834 Å per pixel) with a defocus range from −0.8 to −2.5 µm using SerialEM v4.1.1(*55*). Each image stack was composed of 50 individual frames with a total dose of 50 e^-^ per Å^2^ (for ×105,000 magnification) or 60 e^-^ per Å^2^ (for ×81,000 magnification).

### Cryo-EM image processing

The cryo-EM workflow is shown in **Supplementary Figure 4**. An initial template for particle picking was generated from 4,702 images collected at a magnification of ×81,000 (**Supplementary Figure 4A**). After patch-motion correction and patch contrast transfer function (CTF) estimation in cryoSPARC v4.2(*56, 57*), particles were picked using crYOLO v1.8.4(*58*), extracted with a box size of 256 pixels (1.1 Å/pixel), and binned to 64 pixels in cryoSPARC v4.2. After several rounds of 2D classification and ab initio reconstruction (5 classes), a volume representing the FEX-CA/10E8v4 complex was selected as a template for automatic particle picking for 21,384 patch-motion corrected and CTF-corrected images combined from 3 datasets (×81,000). After many rounds of iterative 2D classification with extracted particles, 938,364 particles were selected and subjected to four rounds of heterogeneous refinement with the previously generated templates, followed by a 2-class ab initio reconstruction. After ab initio reconstruction, 142,174 particles were then subjected to a subsequent round of 3D classification with a solvent mask, and 54,516 particles were selected and refined to 7.28Å resolution using non-uniform refinement. This map showed clear density for the transmembrane helices, but the resolution was insufficient to build an atomic model. We used this low-resolution map to generate 3D template volumes for a FEX-CA/Fab complex, Fab alone, FEX-CA with no Fab, and empty micelles. These four 3D volumes were then used as templates for isolating particles for high-magnification datasets (**Supplementary Figure 4A** – blue box).

An additional 41,056 images were collected at a magnification of ×105,000 and pre-processed using patch-motion correction and patch CTF correction in cryoSPARC v4.5.1 (**Supplementary Figure 4B**). After micrograph denoising, 36,508 images were subjected to templating picking using the low-resolution map of the FEX-CA/Fab complex. Particles were extracted with a box size of 300 pixels (0.834 Å/pixel) followed by heterogeneous refinement with the four 3D template volumes generated earlier (**Supplementary Figure 4A** – blue box). The class representing the Fab/FEX complex was subjected to 4-class ab initio reconstruction followed by heterogeneous refinement. After 3-4 rounds of 2-class ab initio reconstruction followed by heterogenous refinement, a maximal number of complex particles had been obtained. These 439,352 particles were further processed using 2-class ab initio reconstruction followed by 4 rounds of heterogeneous refinement yielding a well-resolved 4.14Å map of the complex with 280,006 particles. To further improve the map resolution, these particles were re-extracted with a larger box size of 352 pixels (0.834 Å/pixel) for iterative 2D classification, 2-class ab initio reconstruction, heterogeneous refinement, and non-uniform refinement, yielding a final 4.0 Å map derived from 164,326 particles (**Supplementary Figure 4C**).

### Model building, refinement, and structural validation

The 10E8v4 Fab structure (PDB: 5IQ9)(*37*) and the FEX-CA Alphafold2 (*59*) prediction were used as initial models to fit into the experimental maps followed by manual adjustment in Coot (v0.9.8.93)(*60*) and real-space refinement in PHENIX (v1.21)(*61*). Structural validation was performed using PHENIX and MolProbity(*62*). PyMOL v3.1 (Schrödinger) and ChimeraX v1.8(*63*) were used to visualize the structural data and generate figures.

### Fluoride efflux assay

Fluoride efflux from proteoliposomes was measured using a lanthanum fluoride electrode (Cole-Parmer, Vernon Hills, IL) as previously described in detail (*12, 64*). Briefly, proteoliposomes were passed over Sephadex G50 resin equilibrated in assay buffer (typically 300 mM K^+^ isethionate, 10 mM HEPES-KOH, pH 7.5) and diluted 40-fold in assay buffer. Fluoride efflux was initiated by the addition of valinomycin (Sigma-Aldrich) to a final concentration of 1.8 µM. At the end of the experiment, proteoliposomes were disrupted by addition of 30 mM n-octyl-β-D-glucoside (β-OG) (Anatrace, Maumee, OH) to release remaining encapsulated fluoride. To convert the fraction of occupied liposomes to the fraction of active protein A_f_, a correction was applied assuming that proteins were inserted into liposomes according to Poisson statistics(*44*):

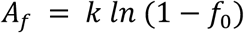

where f_0_ represents the fraction of liposomes that do not exhibit protein-dependent efflux, and *k* is a constant that represents the maximum amount of protein-dependent efflux, which we determine empirically at saturating Na^+^.

### Planar lipid bilayer electrophysiology

Planar lipid bilayer electrophysiology was performed using a horizontal planar lipid bilayer system separated by a partition with *cis* side (upper chamber) and *trans* side (lower chamber, electrical ground) as previously described(*12, 65*). For single channel recording, FEX-CS or FEX-AT proteoliposomes are fused with a 3:1 1-palmitoyl-2-oleoyl-phosphatidylethanolamine (POPE)/ 1-palmitoyl-2-oleoyl-phosphatidylglycerol (POPG) (Avanti Polar Lipids, Alabaster, AL) bilayer, with 300 mM NaF, 15 mM 3-(N-morpholino)propanesulfonic (MOPS) pH 7 in the *cis* chamber, and 30-300 mM NaF, 15 mM MOPS pH 7 in the *trans* chamber, respectively. Current output from an Axopatch 200B amplifier was recorded using Clampex and electronically low-pass filtered at 1 kHz and digitally filtered at 50 Hz for data analysis in Clampfit 10.6 (Molecular Devices, San Jose, CA).

### Fluoride resistance in yeast

The *Δfex1Δfex2* yeast strain SSY3(*7*) was transformed with FEX-CA mutant plasmids using a standard lithium acetate protocol. Transformants were selected on plates prepared with uracil dropout medium (SC-Ura, Sunrise Science Products, Knoxville, TN) with 2% glucose. For yeast resistance assays, overnight cultures were diluted to an OD_600_ of 1 and 10-fold serial dilutions were spotted on SC-Ura plates with 2% galactose to induce FEX expression and NaF as indicated. Plates were imaged after 2-3 days at 30 °C. Expression and membrane localization were validated by Western blot. Cells were collected after 24 hr galactose induction and lysed using Zymolase treatment followed by vortexing. After removing cell debris, the membrane fraction was isolated by ultracentrifugation and membrane pellets were resuspended in buffer containing 4% SDS and 4 M urea. MPER-tagged FEX was detected using purified VRC42 IgG primary antibody(*54*). As a loading control, the constitutively expressed plasma membrane H^+^-ATPase (Pma1) was detected using an anti-Pma1 primary antibody (Invitrogen).

### Molecular dynamic simulations and analysis

The starting structure of FEX-CA was generated using the newly determined cryo-EM structure (PDB: 9OP1). This structure was aligned with Fluc-Bpe (PDB: 5NKQ) to place the Na^+^ ion in the niche. Afterwards, the protonation states of all titratable residues at pH 7.4 were determined using the H++ web server(*66, 67*). The protein was inserted into a 3:1 POPE/POPG bilayer and solvated with TIP3P(*68*) water molecules using Packmol-Memgen(*69*). Missing hydrogen atoms and Na^+^ ions were added to neutralize the system, and NaCl ions were then added using the tleap module from the AmberTools20(*70*) to achieve a physiological salt concentration of 0.150 M. The system consisted of 98,092 atoms. The initial coordinate and parameter files were generated using tleap(*70*), applying the Amber ff14SB(*71*), lipid 21(*72*), and TIP3P(*68*) forcefields.

Three independent simulation trials were performed using the pmemd.cuda module of Amber20 software package(*69, 73*) to ensure adequate sampling. To properly relax and minimize the system, we used a multistep energy minimization protocol in which each step involved both the steepest descent and conjugate gradient methods. Specifically, the first two steps consisted of 5,000 iterations with steepest descent followed by 5,000 iterations with conjugate gradient. In the final step, each method was applied for 10,000 iterations. Initially, only water molecules and hydrogen atoms were minimized, while a positional restraint of 7.5 kcal/mol/Å² was applied to all other atoms. In the second stage, minimization was extended to include ions and protein side chains, with the restraint on the protein backbone. In the final step, all restraints were removed, allowing the entire system to relax freely.

Following minimization, the system was heated in two steps. First, the temperature was gradually increased from 0 to 100 K over 4 ns using the Langevin thermostat(*74*), with restraints of 5 kcal/mol/Å² applied to the protein and lipids. In the second step, the temperature was raised from 100 to 303 K over 8 ns. During this stage, the Berendsen weak-coupling barostat(*75*) was used alongside the Langevin thermostat to equilibrate the pressure, while maintaining the same positional restraints. The system was then equilibrated through 10 sequential steps, each lasting 0.5 ns, to stabilize the periodic boundary conditions. Following equilibration, 750 ns of production simulations were performed for each trial. A 2 fs time step was used, and all bonds involving hydrogen atoms were constrained using the SHAKE algorithm(*76*). A 12 Å cutoff was applied for long-range nonbonded interactions. Simulations were conducted in the NPT ensemble, with the temperature maintained at 303 K using the Langevin thermostat(*74*) and pressure controlled by the Berendsen barostat(*75*). The production frames were saved every 10 ps.

All analyses were performed using the cpptraj module from the AmberTools20(*77*) and VMD software(*78*) on every 100ps of all production trajectories. Root mean square displacement (RMSD) and root mean square fluctuation (RMSF) were calculated to evaluate the structural stability. The number of water molecules within 3 Å and 7 Å of the Na^+^ ion was counted, corresponding to its first and extended coordination shells(*79*). Electrostatic and van der Waals (vdW) interaction energies between the cation-binding motifs and the Na^+^ ion, between the Na^+^ ion and all water molecules within 5 Å, were computed using linear integration energy (lie) analysis. Visualization of the simulations and structural data, as well as figure generation, was performed using VMD software(*78*).

## Acknowledgements

This work was supported by National Institutes of Health grants R35 GM128768 (NIH/NIGMS) to R.B.S., S10OD030275 to M.D.O., and R35GM155106 to H.T. C.-Y. K. was supported by American Heart Association predoctoral fellowship 24PRE1192512. We thank the U-M Cryo-EM facility, which is supported by the U-M Life Sciences Institute, the U-M Biosciences Initiative, and the Beckman Foundation, for scientific and technical assistance, as well as the Office of Information Technology and the Cyber Infrastructure Research Computing (CIRC) at the University of Texas at Dallas and the Texas Advanced Computing Center (TACC) at the University of Texas at Austin for providing HPC resources. We also thank Scott Strobel for the gift of the Δ*fex1*Δ*fex2* yeast strain.

## Author Contributions

C.-Y. K.: conceptualization, investigation, writing – original draft, visualization; M. A.: investigation; S. H.: conceptualization, investigation, data visualization, writing – review and editing; H.T.: conceptualization, writing – review and editing, supervision; M. D. O.: conceptualization, writing – review and editing, supervision; R. B. S.: conceptualization, data visualization, writing – review and editing, supervision, funding acquisition, project administration. The authors declare no competing interests.

## Data and Materials Availability

Coordinates and maps have been deposited in EMDB with PDB ID 9OP1 and EMD-70677. All other data needed to evaluate the conclusions in the paper are present in the paper or the supplementary materials. FEX expression plasmids are available upon request.

---

**Supplementary Figure 1.**
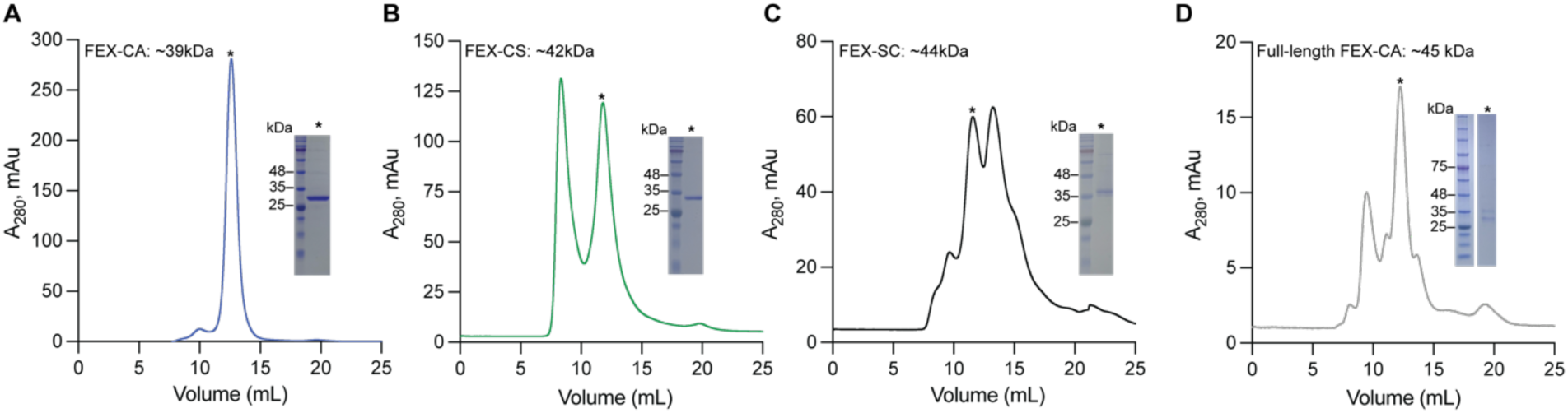
Purification of FEX homologues. Size exclusion chromatograms for FEX homologues from *Candida albicans* (MPER-tagged FEX_Δ1-75_: in panel (A) and full-length FEX in panel (D), *Camellia sinensis* (B), and *Saccharomyces cerevisiae* (C). For each, the inset shows Coomassie-stained SDS-PAGE of peak fractions, indicated by asterisks.

**Supplementary Figure 2.**
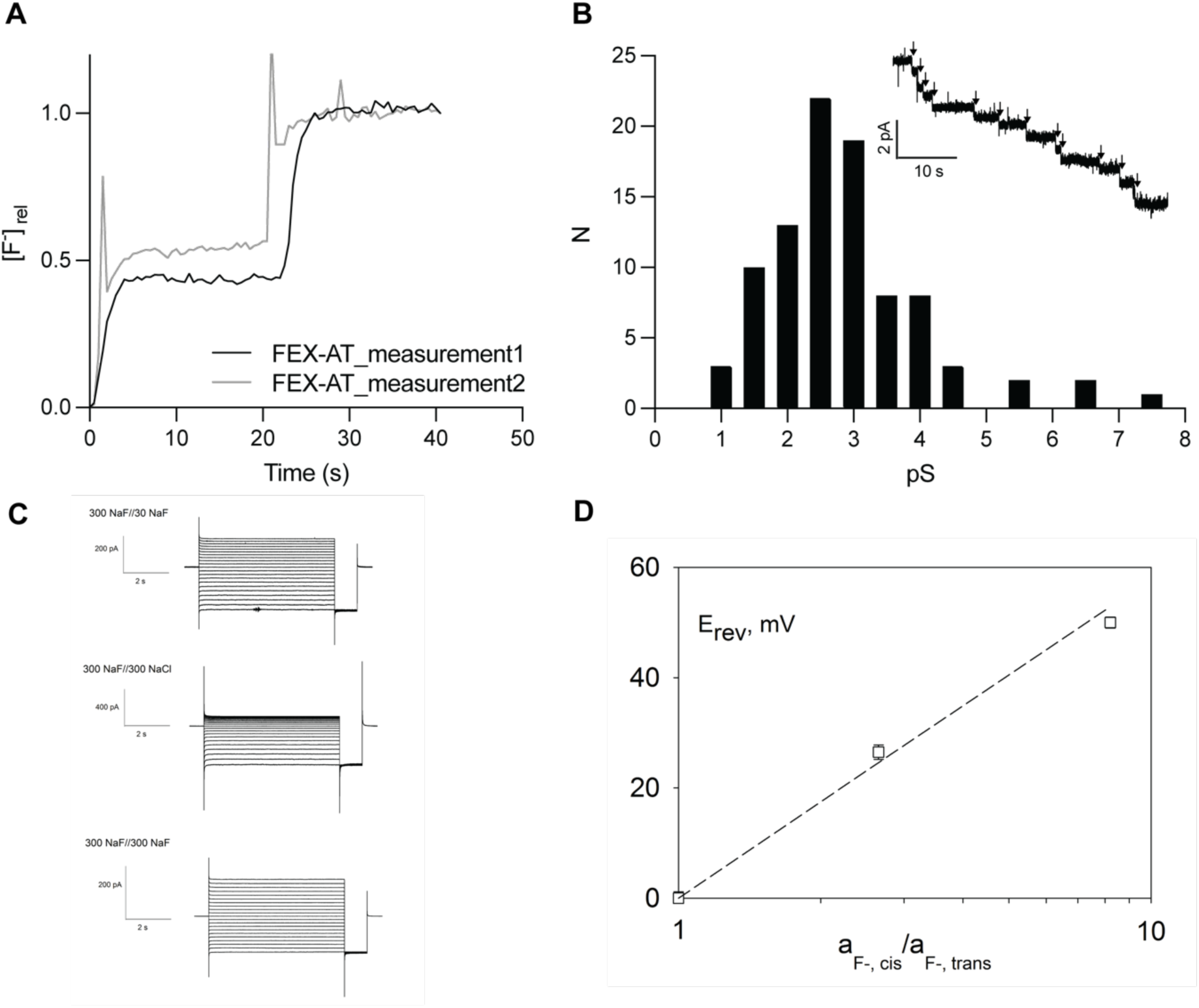
Functional characterization of *Arabidopsis thaliana* FEX. (A) Fluoride efflux from proteoliposomes reconstituted with FEX-AT measured with a fluoride-selective electrode. The reaction was initiated by the addition of valinomycin (first step), and the trace was normalized to the total encapsulated F^-^ determined after detergent addition (second step). (B) Histogram of single channel insertions of FEX-AT. The inset shows a representative electrophysiological recording, with individual FEX-AT insertions into a 3:1 POPE/ POPG lipid bilayer at −200 mV under F^-^ gradient conditions (300 mM NaF // 30 mM NaF) indicated by arrows. (C) Raw current responses to 10-s command voltage pulses (−200 to +200 mV in 20 mV increments) from a holding potential of 0 mV, followed by a 500-ms tail pulse to - 200 mV. The fluoride ion concentration gradient for each (cis//trans) is indicated. (D) Reversal potential as a function of F^-^ activity (a_F_) from three independent bilayers. Points represent the mean and SEM (where not visible, error bars are smaller than the diameter of the point). The dashed line represents the expected relationship between V_r_ and the activity gradient for a selective, electrodiffusive fluoride ion channel. F^-^ activity is calculated from tabulated mean ionic activity coefficients of NaF solutions.

**Supplementary Figure 3.**
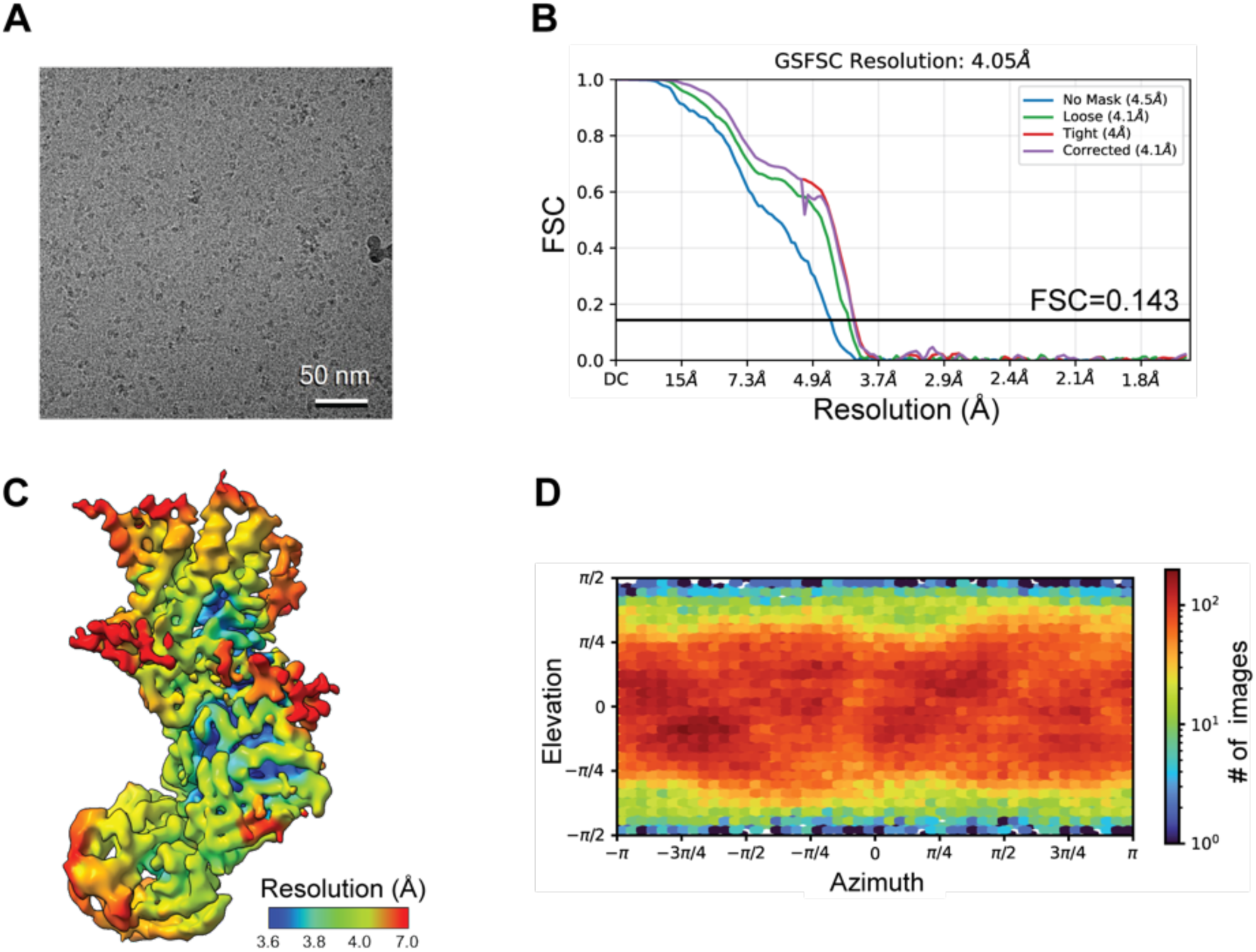
Micrograph and structure validation of FEX. (A) Representative cryo-EM micrograph of FEX-CA in complex with 10E8v4 Fab. (B) FSC curves as a function of resolution calculated between the half map used for refinement and the other half map. (C) Local resolution map colored by local resolution estimate (B factor −181.0) using cryoSPARC. (D) Particle angular distribution plot calculated using cryoSPARC.

**Supplementary Figure 4.**
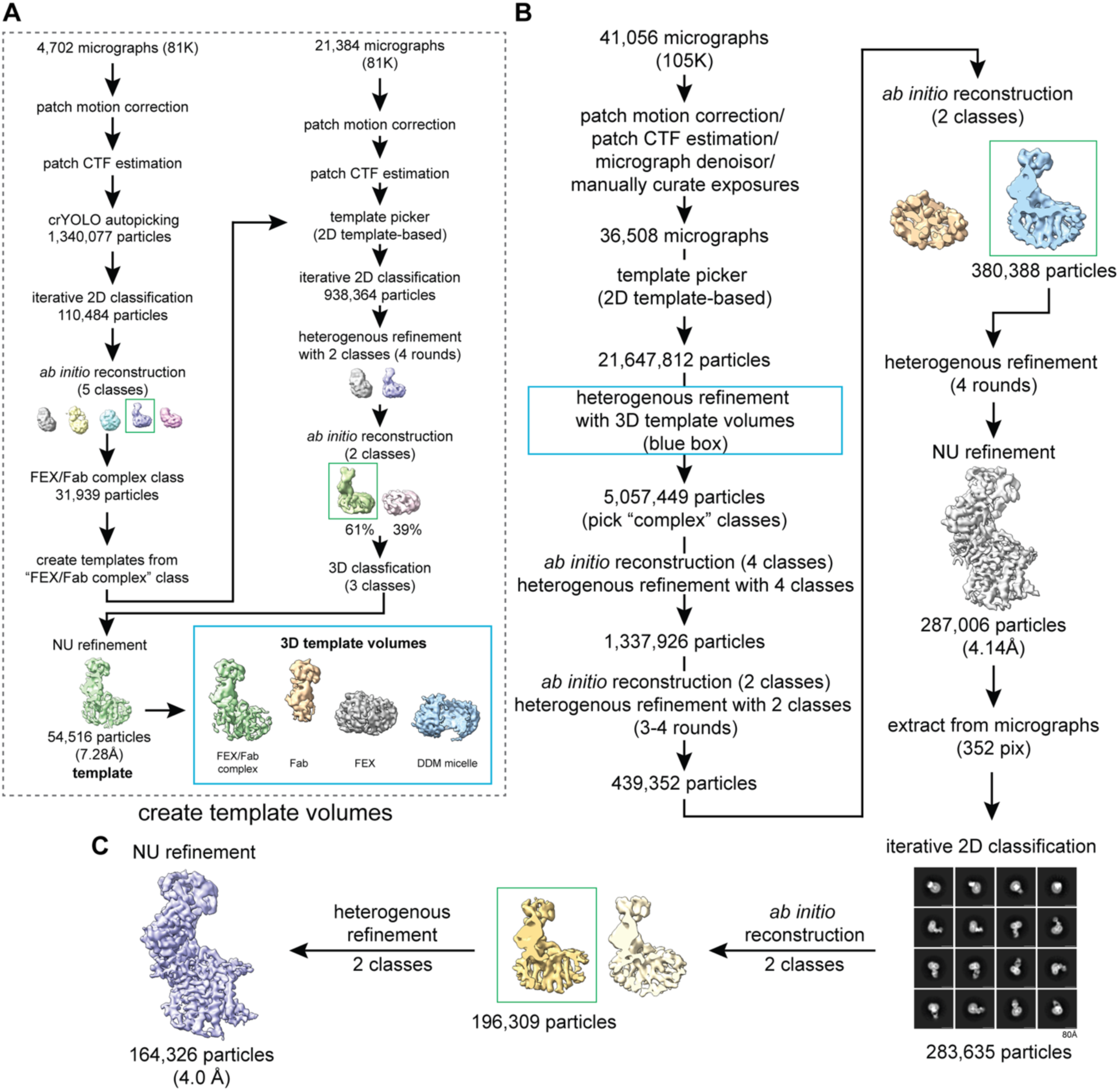
Cryo-EM workflow for FEX-CA data processing. All steps were processed in crYOLO 1.9.2 and Cryosparc v4. 3D template volumes (blue box) were generated from 81,000x magnification dataset (panel A) and used for particle isolation in the 105,000x magnification dataset (panel B). A final class was selected and refined using non-uniform refinement to generate the final 4.0 Å map from 164,326 particles.

**Supplementary Figure 5.**
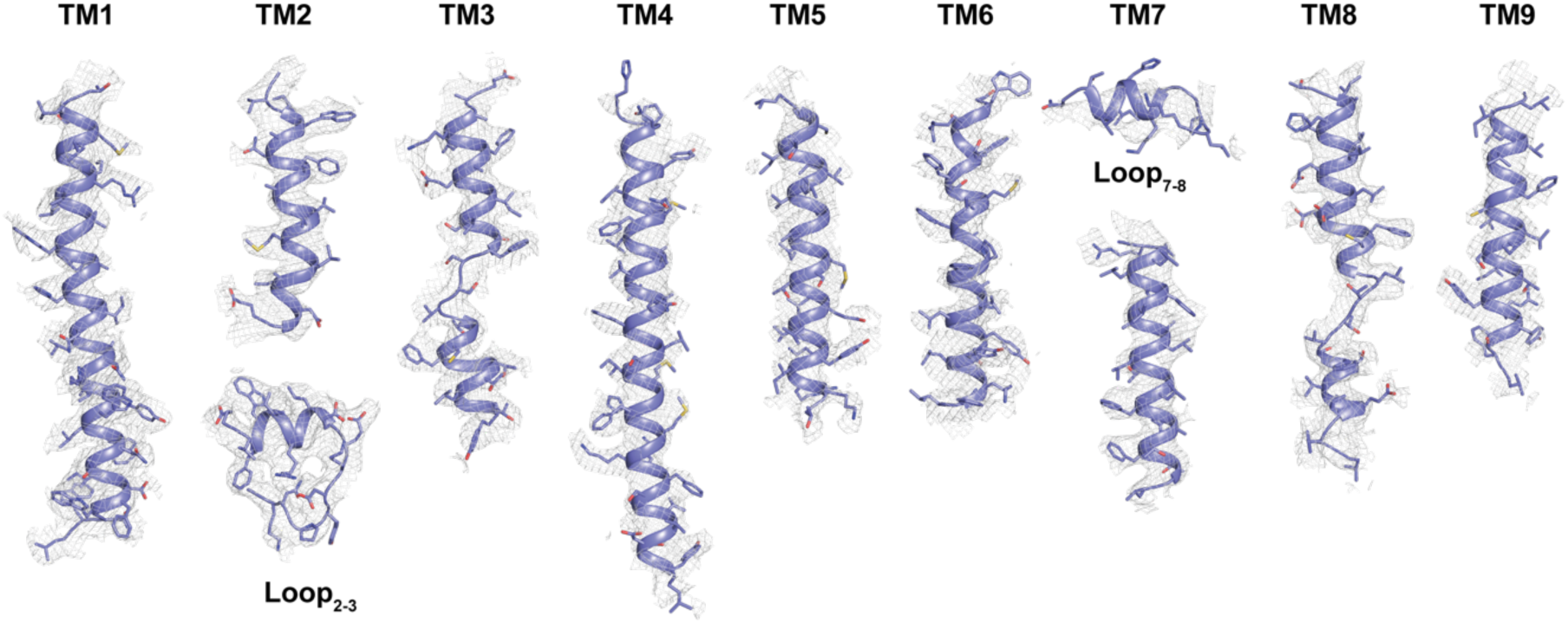
Cryo-EM maps and models of transmembrane helices for FEX-CA. Map contoured at 7σ.

**Supplementary Figure 6.**
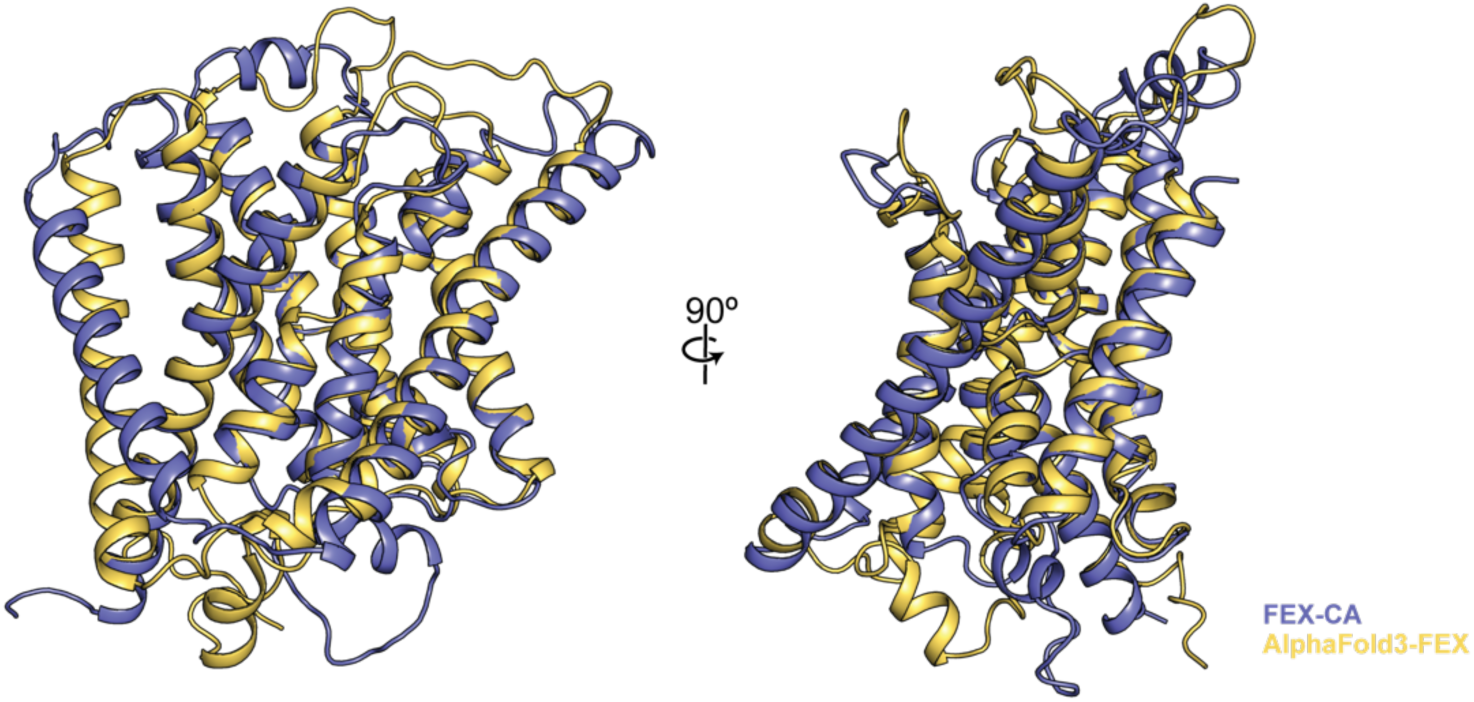
Structure comparison of FEX-CA with model predicted by AlphaFold 3. The experimental FEX-CA model is represented in purple, and the prediction is shown in yellow.

**Supplementary Figure 7.**
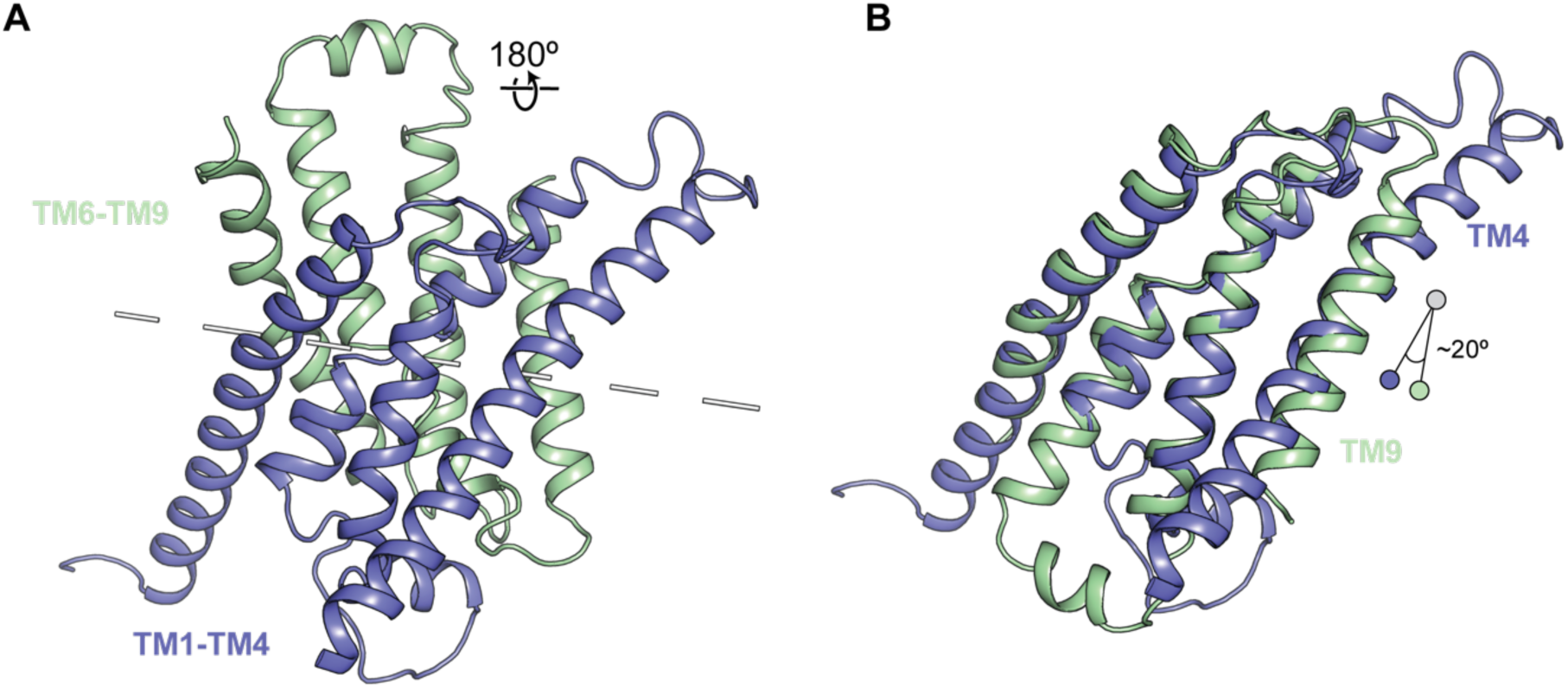
Structure alignment of TM 1-4 and TM 6-9. (A) Pseudosymmetrical domains of TM1-4 and TM 6-9 colored in purple and green, respectively. The internal pseudosymmetry axis is indicated by the dashed line. (B) Overlay of the N- and C-terminal domains.

**Supplementary Figure 8.**
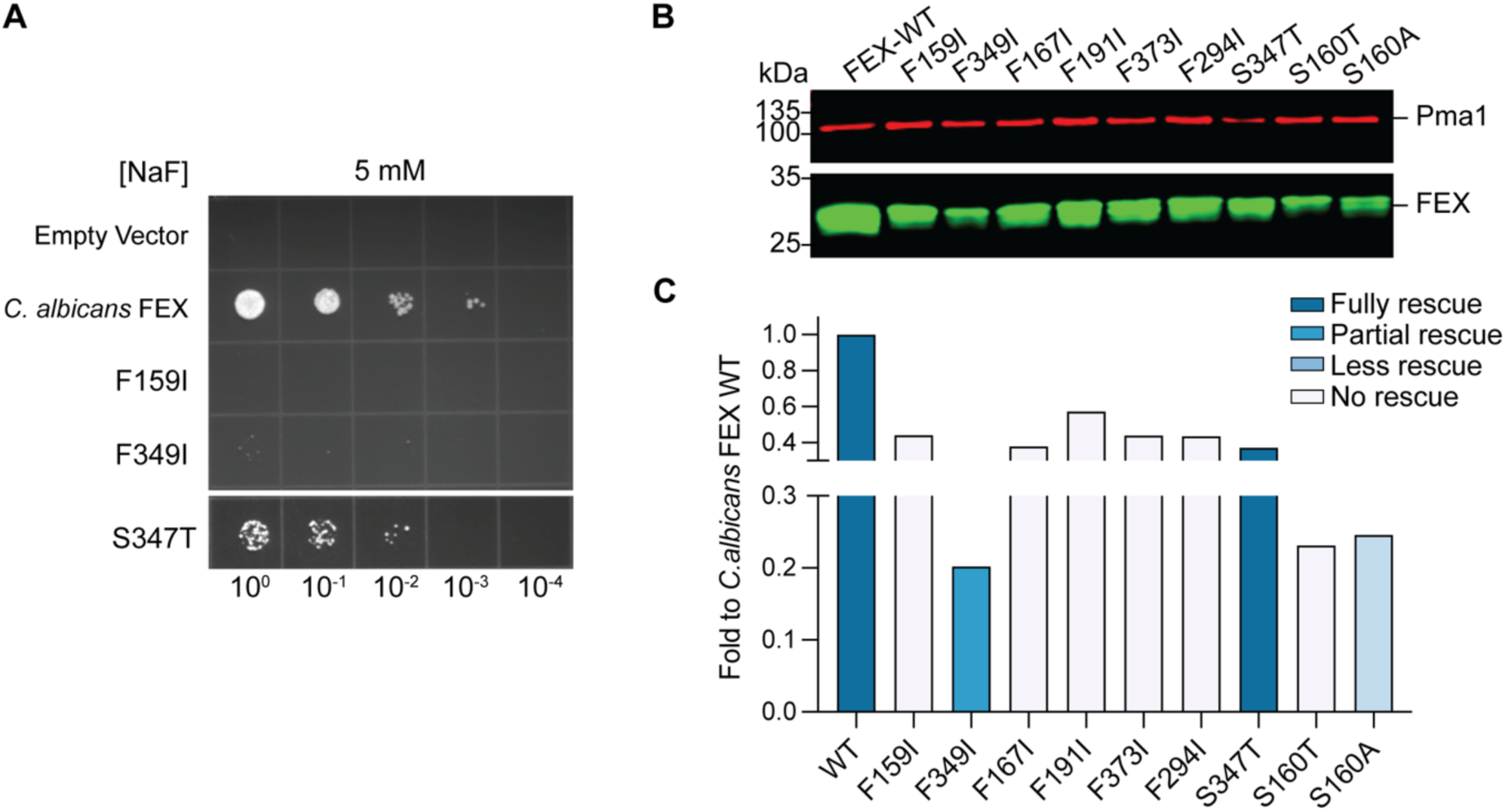
Fluoride resistance and membrane expression of FEX mutants. (A) Ten-fold serial dilutions of Δ*fex1*Δ *fex2 S. cerevisiae* expressing the indicated FEX constructs on -Ura plates with 5 mM NaF. Experiments are representative of 3 independent biological replicates. (B) Western blot analysis of the membrane fraction of Δ*fex1*Δ *fex2 S. cerevisiae* expressing the indicated FEX-CA constructs. Protein is detected using an anti-MPER VRC42 primary antibody and anti-Pma1 antibody as a loading control. (C) Quantification of Western blot band intensity reported as fold change with respect to WT FEX-CA and colored according to the yeast fluoride resistance experiment shown in Fig. 4B.

**Supplementary Figure 9.**
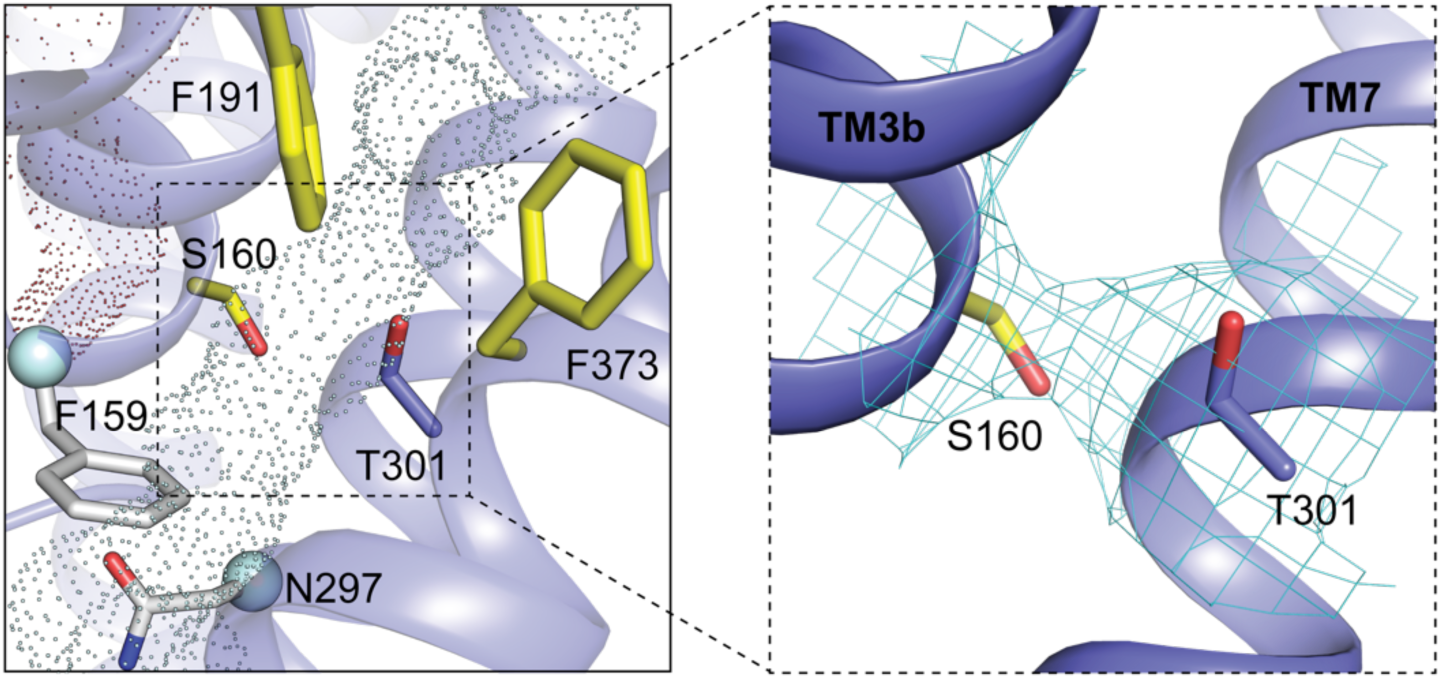
Map of vestibule serine S160 interacting with T301. The zoomed-in view of the predicted permeation path, as shown in Figure 4A, with T301 shown in purple stick representation, with the map surrounding S160 and T301 contoured at 7σ.

**Supplementary Figure 10.**
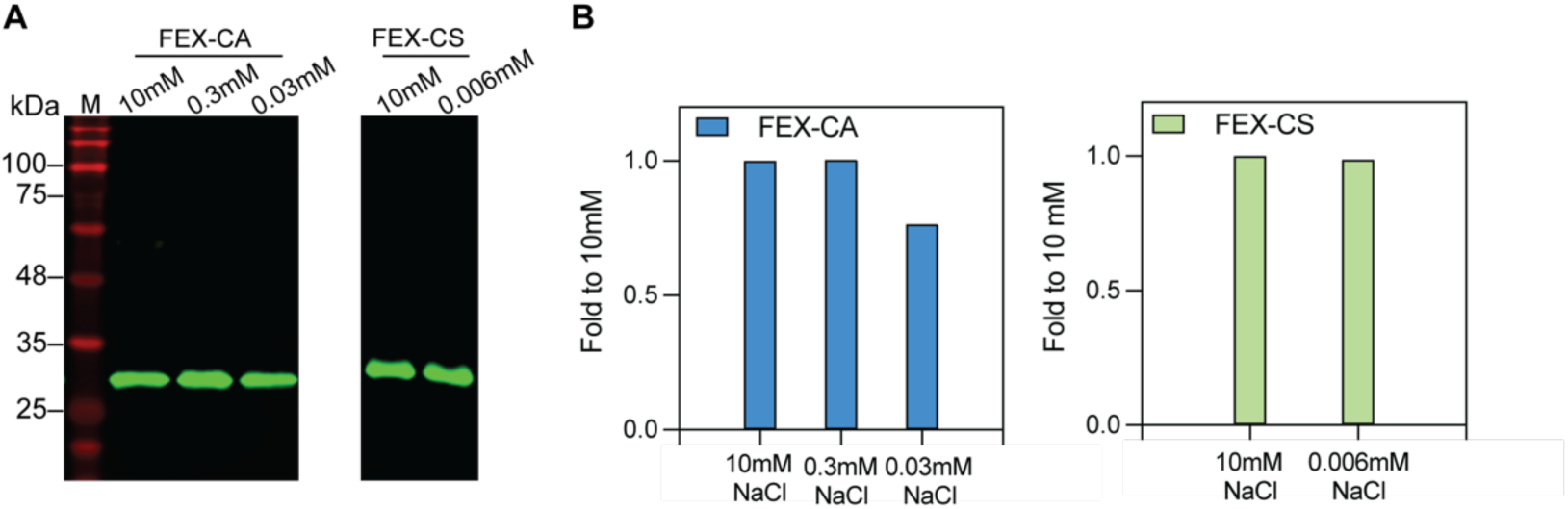
Protein content of proteoliposomes at different concentrations of Na^+^. **(A)** Proteins present in samples from selected high-Na^+^ and low-Na^+^ conditions of the fluoride efflux assay (Figure 5) were detected using anti-MPER antibody. (B) Densiometric quantification of FEX detected in the samples from panel A.

**Supplementary Figure 11.**
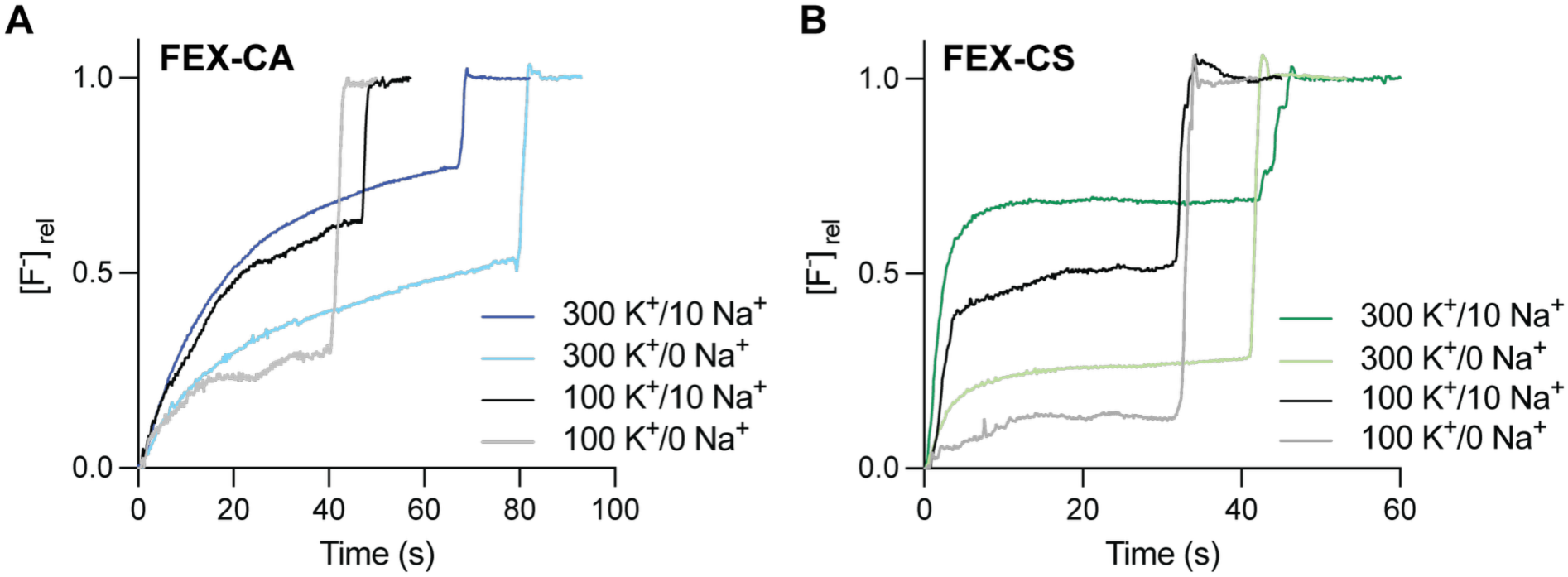
Fluoride efflux activity at different concentrations of K^+^. Fluoride efflux from proteoliposomes reconstituted with FEX-CA (left) and FEX-CS (right) in 100 mM KF or 300 mM KF and Na^+^ as indicated. Traces are representative of three independent experiments. Electrode reading stability decreases at low salt concentrations.

**Supplementary Figure 12.**
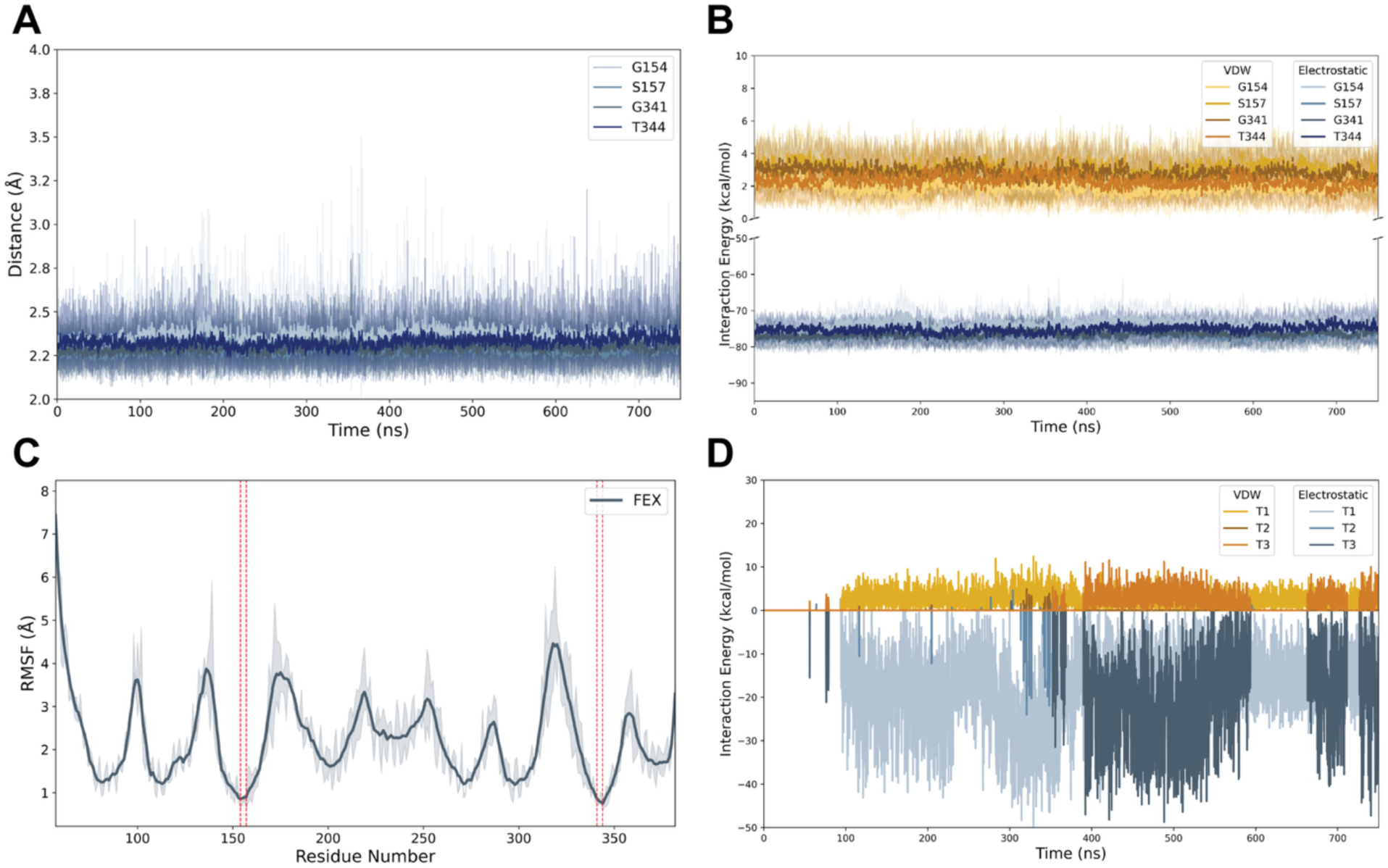
Interaction energy, RMSF, and coordination distance from molecular dynamics simulations. (A) The distance between binding site carbonyl oxygens and the Na^+^ as a function of simulation time, represented as the average (solid line) and s.d (shaded area) of three individual replicates. (B) The interaction energies between the niche carbonyl oxygens and the Na^+^ ion represented as the average (solid lines) and s.d. (shaded area) of three individual replicates. (C) Mean RMSF as a function of FEX residue number shown as the average (solid line) and s.d. (shaded area) of three individual replicates. The Na^+^ coordinating residues G154, S157, G341, and T344 are indicated by the red dashed lines. (D) Calculated interaction energies between the Na^+^ ion and all waters within 3 Å of Na^+^ for each of the three replicates.

**Supplementary Figure 13.**
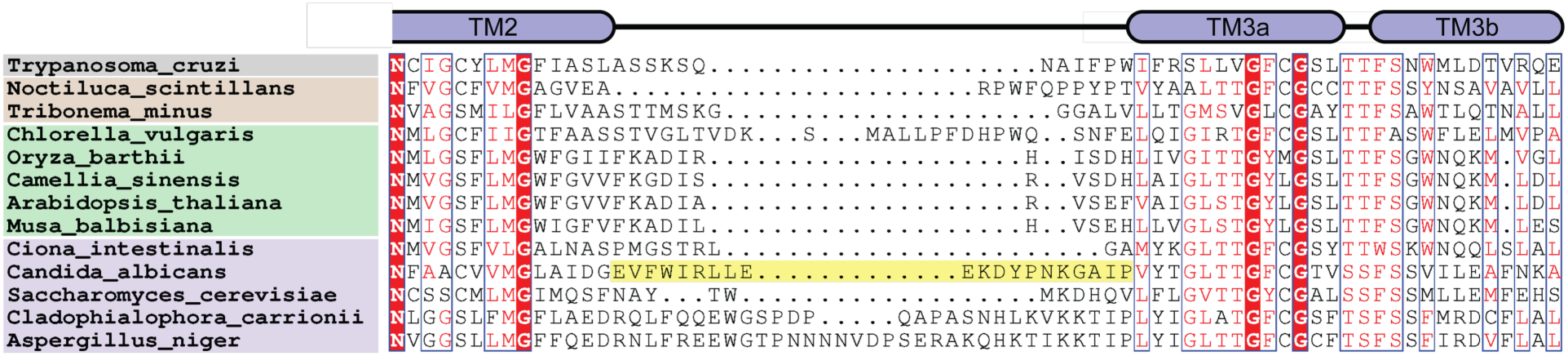
Sequence alignment of TM2/3 linker from thirteen eukaryotic FEX proteins. The alignments were color-coded based on sequence similarity using ESPript 3.0(*83*). Each species is colored according to clade as in Figure 1A. The structured linker between TM2 and TM3 in FEX-CA is highlighted in yellow.

**Supplementary Table 1.** Cryo-EM data collection, refinement, and validation statistics.

|  | FEX-CA<br>(EMD-70677)<br>(PDB 9OP1) |
| --- | --- |
| **Data collection and processing** |  |
| Magnification | 105 K |
| Voltage (kV) | 300 |
| Electron exposure (e-/Å<sup>2</sup>) | 50 |
| Defocus range (μm) | -0.8 to -2.5 |
| Pixel size (Å) | 0.834 |
| Symmetry imposed | C1 symmetry |
| Initial particle images (no.) |  |
| Final particle images (no.) | 164,326 |
| Map resolution (Å) | 4.05 |
| Map resolution range (Å) | 3.5 to 9.1 |
| FSC threshold | 0.143 |
| **Refinement** |  |
| Initial model used | AlphaFold2-predicted model |
| Model resolution (Å) | 3.97 |
| FSC threshold | 0.143 |
| Map sharpening B factor (Å<sup>2</sup>) | 181.0 |
| Model-to-map, CC<sub>mask</sub> | 0.8 |
| Model composition |  |
| Non-hydrogen atoms | 5,843 |
| Protein residues | 766 |
| Water | - |
| B factor (Å<sup>2</sup>) |  |
| Protein | 96.56 |
| Water | - |
| R.m.s. deviations |  |
| Bond lengths (Å) | 0.003 |
| Bond angles (°) | 0.650 |
| **Validation** |  |
| MolProbity score | 2.16 |
| Clashscore | 13.66 |
| Poor rotamers (%) | 0 |
| Ramachandran plot |  |
| Favored (%) | 91.18 |
| Allowed (%) | 8.68 |
| Outliers (%) | 0.13 |

**Supplementary Table 2.**
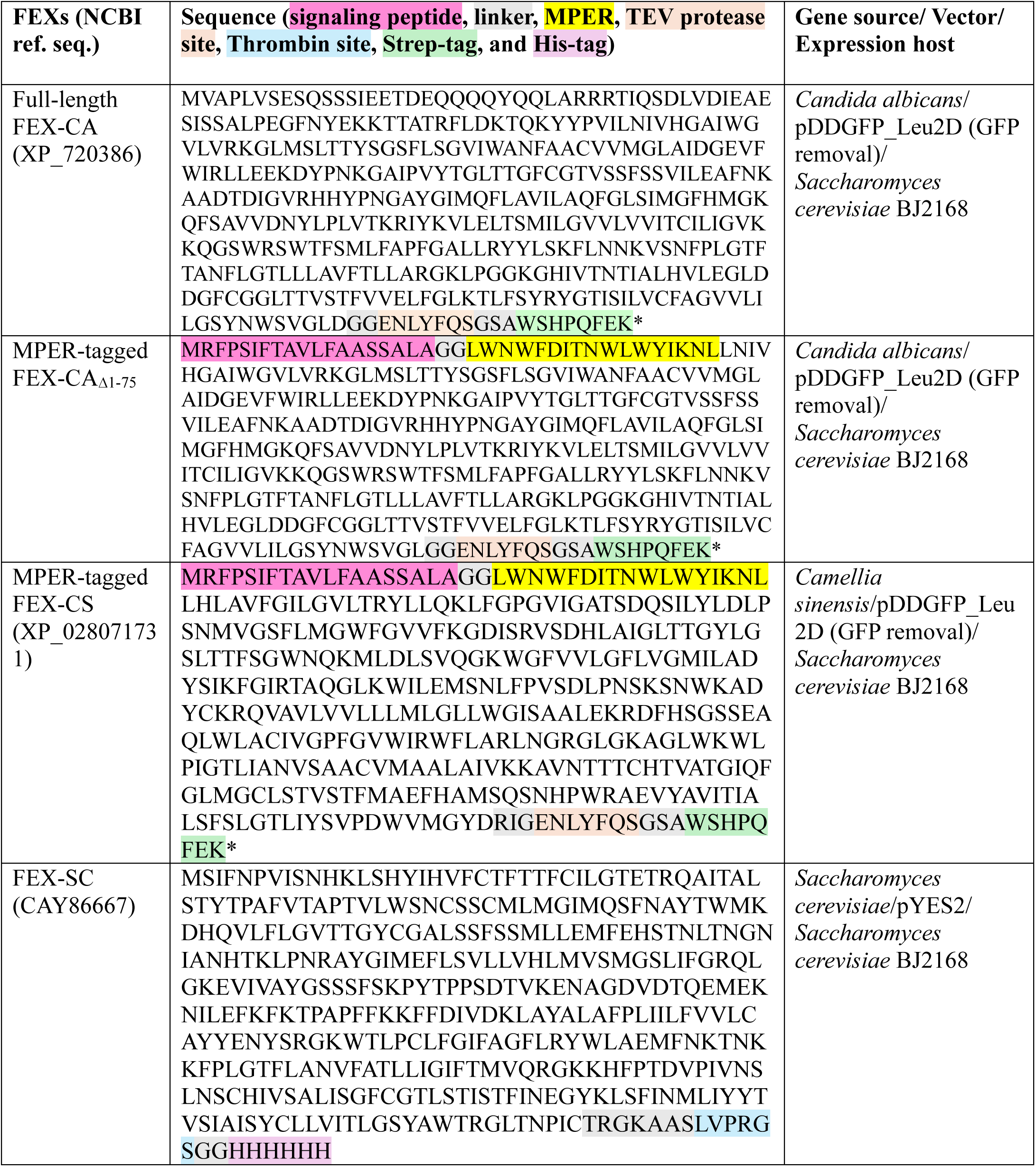
Sequences of FEX homologous constructs.

**Supplementary Table 3:**
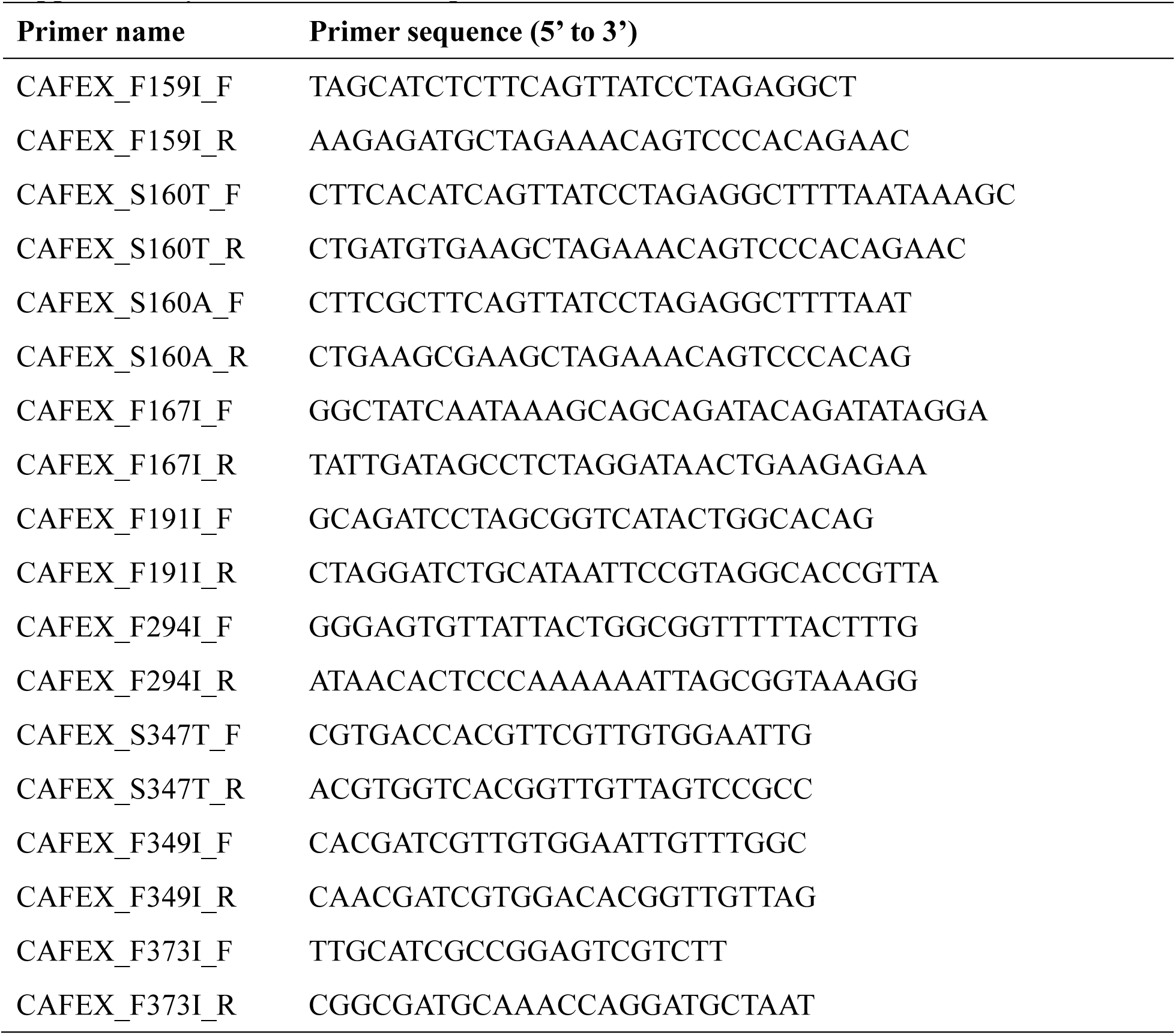
Primer sequences.

## References

1. L. H. Weinstein, A. Davison, Fluorides in the environment : effects on plants and animals. (CABI Pub., Wallingford, Oxon, UK; Cambridge, MA, USA, 2004), pp. ix, 287 p.

2. R. B. Stockbridge, L. P. Wackett, The link between ancient microbial fluoride resistance mechanisms and bioengineering organofluorine degradation or synthesis. Nature Communications 15, 4593 (2024).

3. V. R. Samygina et al., Reversible inhibition of Escherichia coli inorganic pyrophosphatase by fluoride: trapped catalytic intermediates in cryo-crystallographic studies. J Mol Biol 366, 1305–1317 (2007).

4. O. Barbier, L. Arreola-Mendoza, L. M. Del Razo, Molecular mechanisms of fluoride toxicity. Chem Biol Interact 188, 319–333 (2010).

5. E. Adamek, K. Pawlowska-Goral, K. Bober, In vitro and in vivo effects of fluoride ions on enzyme activity. Ann Acad Med Stetin 51, 69–85 (2005).

6. J. L. Baker et al., Widespread genetic switches and toxicity resistance proteins for fluoride. Science 335, 233–235 (2012).

7. S. Li et al., Eukaryotic resistance to fluoride toxicity mediated by a widespread family of fluoride export proteins. Proceedings of the National Academy of Sciences 110, 19018–19023 (2013).

8. A. Banerjee, A. Roychoudhury, Functional and molecular characterization of fluoride exporter (FEX) from rice and its constitutive overexpression in Nicotiana benthamiana to promote fluoride tolerance. Plant Cell Rep 40, 1751–1772 (2021).

9. A. Banerjee et al., Fluoride export is required for competitive fitness of pathogenic microorganisms in dental biofilm models. mBio 0:e00184–24, (2024).

10. B. C. McIlwain, M. T. Ruprecht, R. B. Stockbridge, Membrane Exporters of Fluoride Ion. Annu Rev Biochem 90, 559–579 (2021).

11. R. B. Stockbridge et al., Fluoride resistance and transport by riboswitch-controlled CLC antiporters. Proc Natl Acad Sci U S A 109, 15289–15294 (2012).

12. R. B. Stockbridge, J. L. Robertson, L. Kolmakova-Partensky, C. Miller, A family of fluoride-specific ion channels with dual-topology architecture. Elife 2, e01084 (2013).

13. T. Berbasova et al., Fluoride export (FEX) proteins from fungi, plants and animals are ‘single barreled’ channels containing one functional and one vestigial ion pore. PLoS One 12, e0177096 (2017).

14. S. L. Tausta, T. Berbasova, M. Peverelli, S. A. Strobel, The fluoride transporter FLUORIDE EXPORTER (FEX) is the major mechanism of tolerance to fluoride toxicity in plants. Plant Physiol, (2021).

15. R. B. Stockbridge, A. Koide, C. Miller, S. Koide, Proof of dual-topology architecture of Fluc F-channels with monobody blockers. Nat Commun 5, 5120 (2014).

16. K. D. Smith et al., Yeast Fex1p Is a Constitutively Expressed Fluoride Channel with Functional Asymmetry of Its Two Homologous Domains. Journal of Biological Chemistry 290, 19874–19887 (2015).

17. R. Keller, C. Ziegler, D. Schneider, When two turn into one: evolution of membrane transporters from half modules. Biol Chem 395, 1379–1388 (2014).

18. A. A. Aleksandrova, E. Sarti, L. R. Forrest, MemSTATS: A Benchmark Set of Membrane Protein Symmetries and Pseudosymmetries. J Mol Biol 432, 597–604 (2020).

19. C. B. Macdonald, R. B. Stockbridge, A topologically diverse family of fluoride channels. Curr Opin Struct Biol 45, 142–149 (2017).

20. D. L. Turman, J. T. Nathanson, R. B. Stockbridge, T. O. Street, C. Miller, Two-sided block of a dual-topology F-channel. Proc Natl Acad Sci U S A 112, 5697–5701 (2015).

21. D. L. Turman, R. B. Stockbridge, Mechanism of single- and double-sided inhibition of dual topology fluoride channels by synthetic monobodies. J Gen Physiol, (2017).

22. R. B. Stockbridge et al., Crystal structures of a double-barrelled fluoride ion channel. Nature 525, 548–551 (2015).

23. N. B. Last, L. Kolmakova-Partensky, T. Shane, C. Miller, Mechanistic signs of double-barreled structure in a fluoride ion channel. Elife 5, (2016).

24. B. C. McIlwain, R. Gundepudi, B. B. Koff, R. B. Stockbridge, The fluoride permeation pathway and anion recognition in Fluc family fluoride channels. Elife 10, (2021).

25. J. Zhang et al., Fluoride permeation mechanism of the Fluc channel in liposomes revealed by solid-state NMR. Sci Adv 9, eadg9709 (2023).

26. N. B. Last, S. Sun, M. C. Pham, C. Miller, Molecular determinants of permeation in a fluoride-specific ion channel. Elife 6, (2017).

27. Z. Yue, Z. Wang, G. A. Voth, Ion permeation, selectivity, and electronic polarization in fluoride channels. Biophys J 121, 1336–1347 (2022).

28. M. Ernst, E. A. Orabi, R. B. Stockbridge, J. D. Faraldo-Gomez, J. L. Robertson, Dimerization mechanism of an inverted-topology ion channel in membranes. Proc Natl Acad Sci U S A 120, e2308454120 (2023).

29. B. C. McIlwain, K. Martin, E. A. Hayter, R. B. Stockbridge, An Interfacial Sodium Ion is an Essential Structural Feature of Fluc Family Fluoride Channels. J Mol Biol 432, 1098–1108 (2020).

30. K. D. Smith et al., Yeast Fex1p Is a Constitutively Expressed Fluoride Channel with Functional Asymmetry of Its Two Homologous Domains. J Biol Chem 290, 19874–19887 (2015).

31. A. Rivetta, C. Slayman, Electrophysiology of fluoride channels in the yeasts Saccharomyces cerevisiae and Candida albicans. Methods Enzymol 696, 3–24 (2024).

32. J. Song et al., Two New Members of CsFEXs Couple Proton Gradients to Export Fluoride and Participate in Reducing Fluoride Accumulation in Low-Fluoride Tea Cultivars. J Agric Food Chem 68, 8568–8579 (2020).

33. J. Yang et al., Critical review of fluoride in tea plants (Camellia sinensis): absorption, transportation, tolerance mechanisms, and defluorination measures. Beverage Plant Research 4, (2024).

34. N. I. Nicely et al., Crystal structure of a non-neutralizing antibody to the HIV-1 gp41 membrane-proximal external region. Nat Struct Mol Biol 17, 1492–1494 (2010).

35. J. Huang et al., Broad and potent neutralization of HIV-1 by a gp41-specific human antibody. Nature 491, 406–412 (2012).

36. C. Soto et al., Developmental Pathway of the MPER-Directed HIV-1-Neutralizing Antibody 10E8. PLoS One 11, e0157409 (2016).

37. Y. D. Kwon et al., Optimization of the Solubility of HIV-1-Neutralizing Antibody 10E8 through Somatic Variation and Structure-Based Design. J Virol 90, 5899–5914 (2016).

38. B. C. McIlwain et al., N-terminal Transmembrane-Helix Epitope Tag for X-ray Crystallography and Electron Microscopy of Small Membrane Proteins. J Mol Biol 433, 166909 (2021).

39. P. Li et al., Substrate transport and drug interaction of human thiamine transporters SLC19A2/A3. Nat Commun 15, 10924 (2024).

40. B. C. McIlwain, S. Newstead, R. B. Stockbridge, Cork-in-Bottle Occlusion of Fluoride Ion Channels by Crystallization Chaperones. Structure 26, 635–639 e631 (2018).

41. J. Abramson et al., Accurate structure prediction of biomolecular interactions with AlphaFold 3. Nature 630, 493–500 (2024).

42. N. B. Last et al., A CLC-type F(-)/H(+) antiporter in ion-swapped conformations. Nat Struct Mol Biol 25, 601–606 (2018).

43. B. C. McIlwain, K. Martin, E. A. Hayter, R. B. Stockbridge, An Interfacial Sodium Ion is an Essential Structural Feature of Fluc Family Fluoride Channels. J Mol Biol, (2020).

44. R. B. Stockbridge, The application of Poisson distribution statistics in ion channel reconstitution to determine oligomeric architecture. Methods Enzymol 652, 321–340 (2021).

45. G. M. Torrance, D. P. Leader, D. R. Gilbert, E. J. Milner-White, A novel main chain motif in proteins bridged by cationic groups: the niche. J Mol Biol 385, 1076–1086 (2009).

46. A. E. Brammer, R. B. Stockbridge, C. Miller, F-/Cl-selectivity in CLCF-type F-/H+ antiporters. J Gen Physiol 144, 129–136 (2014).

47. H. Innan, F. Kondrashov, The evolution of gene duplications: classifying and distinguishing between models. Nat Rev Genet 11, 97–108 (2010).

48. J. Levring, J. Chen, Structural identification of a selectivity filter in CFTR. Proc Natl Acad Sci U S A 121, e2316673121 (2024).

49. K. Khafizov et al., Investigation of the sodium-binding sites in the sodium-coupled betaine transporter BetP. Proc Natl Acad Sci U S A 109, E3035–3044 (2012).

50. P. Hariharan, L. Guan, Thermodynamic cooperativity of cosubstrate binding and cation selectivity of Salmonella typhimurium MelB. J Gen Physiol 149, 1029–1039 (2017).

51. S. G. Schmidt, A. Nygaard, J. A. Mindell, C. J. Loland, Exploring the K(+) binding site and its coupling to transport in the neurotransmitter:sodium symporter LeuT. Elife 12, (2024).

52. G. P. Lin-Cereghino et al., The effect of alpha-mating factor secretion signal mutations on recombinant protein expression in Pichia pastoris. Gene 519, 311–317 (2013).

53. J. L. Parker, S. Newstead, Method to increase the yield of eukaryotic membrane protein expression in Saccharomyces cerevisiae for structural and functional studies. Protein Sci 23, 1309–1314 (2014).

54. S. J. Krebs et al., Longitudinal Analysis Reveals Early Development of Three MPER-Directed Neutralizing Antibody Lineages from an HIV-1-Infected Individual. Immunity 50, 677–691 e613 (2019).

55. D. N. Mastronarde, Automated electron microscope tomography using robust prediction of specimen movements. J Struct Biol 152, 36–51 (2005).

56. A. Punjani, J. L. Rubinstein, D. J. Fleet, M. A. Brubaker, cryoSPARC: algorithms for rapid unsupervised cryo-EM structure determination. Nat Methods 14, 290–296 (2017).

57. A. Punjani, H. Zhang, D. J. Fleet, Non-uniform refinement: adaptive regularization improves single-particle cryo-EM reconstruction. Nat Methods 17, 1214–1221 (2020).

58. T. Wagner et al., SPHIRE-crYOLO is a fast and accurate fully automated particle picker for cryo-EM. Commun Biol 2, 218 (2019).

59. J. Jumper et al., Highly accurate protein structure prediction with AlphaFold. Nature 596, 583-+ (2021).

60. P. Emsley, B. Lohkamp, W. G. Scott, K. Cowtan, Features and development of Coot. Acta Crystallogr D Biol Crystallogr 66, 486–501 (2010).

61. P. V. Afonine et al., Real-space refinement in PHENIX for cryo-EM and crystallography. Acta Crystallogr D Struct Biol 74, 531–544 (2018).

62. M. G. Prisant, C. J. Williams, V. B. Chen, J. S. Richardson, D. C. Richardson, New tools in MolProbity validation: CaBLAM for CryoEM backbone, UnDowser to rethink "waters," and NGL Viewer to recapture online 3D graphics. Protein Sci 29, 315–329 (2020).

63. E. F. Pettersen et al., UCSF ChimeraX: Structure visualization for researchers, educators, and developers. Protein Sci 30, 70–82 (2021).

64. C. Y. Kang, M. An, R. B. Stockbridge, Lanthanum-fluoride electrode-based methods to monitor fluoride transport in cells and reconstituted lipid vesicles. Methods Enzymol 696, 43–63 (2024).

65. A. Accardi, L. Kolmakova-Partensky, C. Williams, C. Miller, Ionic currents mediated by a prokaryotic homologue of CLC Cl-channels. J Gen Physiol 123, 109–119 (2004).

66. R. Anandakrishnan, B. Aguilar, A. V. Onufriev, H++ 3.0: automating pK prediction and the preparation of biomolecular structures for atomistic molecular modeling and simulations. Nucleic Acids Res 40, W537–541 (2012).

67. J. C. Gordon et al., H++: a server for estimating pKas and adding missing hydrogens to macromolecules. Nucleic Acids Res 33, W368–371 (2005).

68. W. L. Jorgensen, J. Chandrasekhar, J. D. Madura, R. W. Impey, M. L. Klein, Comparison of Simple Potential Functions for Simulating Liquid Water. J Chem Phys 79, 926–935 (1983).

69. S. Schott-Verdugo, H. Gohlke, PACKMOL-Memgen: A Simple-To-Use, Generalized Workflow for Membrane-Protein-Lipid-Bilayer System Building. J Chem Inf Model 59, 2522–2528 (2019).

70. D. A. Case et al., AMBER 2020. (University of California, San Francisco, 2020).

71. J. A. Maier et al., ff14SB: Improving the Accuracy of Protein Side Chain and Backbone Parameters from ff99SB. J Chem Theory Comput 11, 3696–3713 (2015).

72. C. J. Dickson, R. C. Walker, I. R. Gould, Lipid21: Complex Lipid Membrane Simulations with AMBER. J Chem Theory Comput 18, 1726–1736 (2022).

73. A. W. Gotz et al., Routine Microsecond Molecular Dynamics Simulations with AMBER on GPUs. 1. Generalized Born. J Chem Theory Comput 8, 1542–1555 (2012).

74. R. J. Loncharich, B. R. Brooks, R. W. Pastor, Langevin dynamics of peptides: the frictional dependence of isomerization rates of N-acetylalanyl-N’-methylamide. Biopolymers 32, 523–535 (1992).

75. H. J. C. Berendsen, J. P. M. Postma, W. F. Vangunsteren, A. Dinola, J. R. Haak, Molecular-Dynamics with Coupling to an External Bath. J Chem Phys 81, 3684–3690 (1984).

76. J. P. Ryckaert, G. Ciccotti, H. J. C. Berendsen, Numerical integration of the cartesian equations of motion of a system with constraints: molecular dynamics of n-alkanes. Journal of computational physics 23, 327–341 (1977).

77. D. R. Roe, T. E. Cheatham, 3rd, PTRAJ and CPPTRAJ: Software for Processing and Analysis of Molecular Dynamics Trajectory Data. J Chem Theory Comput 9, 3084–3095 (2013).

78. W. Humphrey, A. Dalke, K. Schulten, VMD: visual molecular dynamics. J Mol Graph 14, 33-38, 27–38 (1996).

79. H. Yu et al., Simulating Monovalent and Divalent Ions in Aqueous Solution Using a Drude Polarizable Force Field. J Chem Theory Comput 6, 774–786 (2010).

80. F. Burki, A. J. Roger, M. W. Brown, A. G. B. Simpson, The New Tree of Eukaryotes. Trends Ecol Evol 35, 43–55 (2020).

81. C. UniProt, UniProt: the Universal Protein Knowledgebase in 2025. Nucleic Acids Res 53, D609–D617 (2025).

82. E. Chovancova et al., CAVER 3.0: a tool for the analysis of transport pathways in dynamic protein structures. PLoS Comput Biol 8, e1002708 (2012).

83. X. Robert, P. Gouet, Deciphering key features in protein structures with the new ENDscript server. Nucleic Acids Res 42, W320–324 (2014).

